# Dynamic Myosin 10 coupling to DCC and β1 integrin is mediated by intrinsically disordered regions during filopodial transport and patterning

**DOI:** 10.64898/2026.04.10.717811

**Authors:** Julia Shangguan, Susanne Reinhardt, Shao Huan Samuel Weng, Ralf Jungmann, Tobin R. Sosnick, Ronald S. Rock

## Abstract

Intrinsically disordered regions (IDRs) are key mediators of protein-protein interactions. IDRs are important components of Myosin 10 (Myo10) and cargo complexes that influence neuronal development and cell growth, yet how IDRs dictate Myo10’s cargo affinity and selectivity is not fully understood. Here, we investigate how the actin motor protein Myo10 engages two distinct cargo receptors, DCC and β1 integrin, in cellular protrusions known as filopodia. Using hydrogen-deuterium exchange mass spectrometry (HDX-MS), cross-linking mass spectrometry (XL-MS), live-cell imaging, and super-resolution microscopy, we show that Myo10 decodes IDR elements through two complementary mechanisms: disorder-to-order transitions and “fuzzy” binding. The cytoplasmic portion of DCC binds Myo10 via a weakly helical P3 motif that acts as a preformed recognition element, while additional disordered motifs contribute to affinity through dynamic, weak interactions. In contrast, the β1 integrin tail interacts with Myo10 through short NPxY motifs that remain disordered. Both cargos engage a common Myo10 surface but also contact distinct sites. Super-resolution DNA-PAINT imaging reveals distinct patterning of cargo with Myo10 along and around filopodia. Concentration measurements show that DCC is primarily bound while β1 integrin exhibits a broader range of occupancy along the filopodial shaft. Multiple additive weak contacts and a shared binding site implies that DCC can out-compete integrin for Myo10 binding, which causes redistribution of active β1 integrin from the filopodial tip to the shaft. Our findings illustrate a tunable, multivalent binding strategy that allows Myo10 to selectively coordinate diverse signaling cargos, demonstrating how regulated disorder within IDRs is one mechanism underlying cargo binding and cellular signaling.

## Introduction

Myo10 is an unconventional motor that drives many essential signaling processes in both normal and pathological cellular functions.^1,2^ Through selective interactions with actin bundles^3,4^ and cargo proteins, it promotes the formation of filopodia,^5–7^ a class of cellular protrusions that carry out mechano-sensing, migration, and phagocytosis.^8^ As with many other unconventional myosins (i.e., nonmuscle), Myo10 is comprised of three primary domains: a head that binds actin filaments and undergoes ATP hydrolysis; a neck that binds lipids and amplifies movements; and a C-terminal tail that binds cargo. A conserved MyTH4-FERM domain (M4F) in the Myo10 C-terminal tail interacts with a variety of cargo proteins including VASP,^9^ microtubules,^10,11^ bone morphogenetic protein receptor,^12^ vascular-endothelial cadherin,^13^ integrin^11,14^ and Deleted in Colorectal Cancer (DCC)^11,15,16^. Myo10’s cargo-binding ability, particularly with integrin and DCC, directly impacts pathways central to tumor progression and neuronal guidance.

Integrins are transmembrane receptors that adhere and respond to extracellular matrix (ECM) proteins and establish connections to the actin cytoskeleton.^17^ As heterodimeric receptors, they comprise α and β subunits. Depending on the integrin class and splice variant of β cytoplasmic domains, distinct conformational arrangements arise that affect affinity to partners such as talin and filamin;^18^ these interactions are regulated by inside-out signaling.^19^ In the case of integrin subunit β1, Myo10 attaches β1 integrin to filopodial tips for protrusion stability and cell adhesion.^14,20^ Integrin signaling can drive cancer cell outgrowth while the loss of Myo10-integrin binding diminishes cancer cell invasion, suggesting that Myo10 contributes to cancer progression.^21^ Indeed, Myo10 is upregulated in several cancers.^22^

DCC is a single-pass transmembrane receptor that binds the netrin family of extracellular guidance cues to stimulate and steer axon outgrowth. Myo10 transports DCC to neurite tips and is essential for netrin-1-guided axon projection and pathfinding.^23^ Coordination of Myo10 with DCC, netrin-1, and KIF13B (a kinesin family motor protein) allows for axonal distribution and Myo10 transport in neurons, promoting Myo10-regulated axonal initiation and branching.^24^ Through mechanistic studies, Yu et al. found that Myo10 knock-down in embryonic mouse neurons disturbs netrin-1 promoted axon initiation and branching/targeting.^24^ Outside of its neuronal context, DCC also plays a role in cancer, where loss of its expression in colorectal carcinoma tumor cells is associated with lower patient survival.^25^

Because Myo10’s interactions with integrin and DCC are essential to important cellular processes, it is valuable to investigate how Myo10 recognizes its cargo, preferentially selects cargo types, switches between cargoes, and simultaneously coordinates the activity of multiple cargos. Clues regarding how DCC and β1 integrin interact with Myo10 are present in existing structural and biochemical data (Figure 1). Myo10’s M4F domain contacts the cytoplasmic tails of β1 integrin and DCC. The cytoplasmic β1 integrin tail is short (∼ 40 residues) and contains two NPxY motifs that can bind Myo10^14,20^. No crystal structure is available for the integrin tail bound to the Myo10 M4F. However, related structures of β integrin tails bound to the talin FERM domains show that one NPxY motif forms a reverse turn that docks near the FERM C-terminal helix^26,27^. DCC’s cytoplasmic tail is over 300 residues long and is primarily an intrinsically disordered region (IDR). The DCC tail contains short, conserved motifs named P1, P2, and P3,^28^ where the P3 motif forms an α-helix that binds to the C-terminal helix of the Myo10 M4F.^11,15^ A common feature of β1 integrin and DCC is that they both have small M4F-binding motifs, with buried surface areas of 550 and 970 Å^2^, respectively. Thus, an open question is how Myo10 achieves long-range transport down a filopodium while barely contacting its cargo. For DCC at least, its IDRs may contribute through “fuzzy” binding^29^ or disorder-to-order transitions.^30–32^ Despite their presence in DCC and integrin tails, IDRs have been traditionally ignored in studies of Myo10 cargo binding.

**Figure 1.**
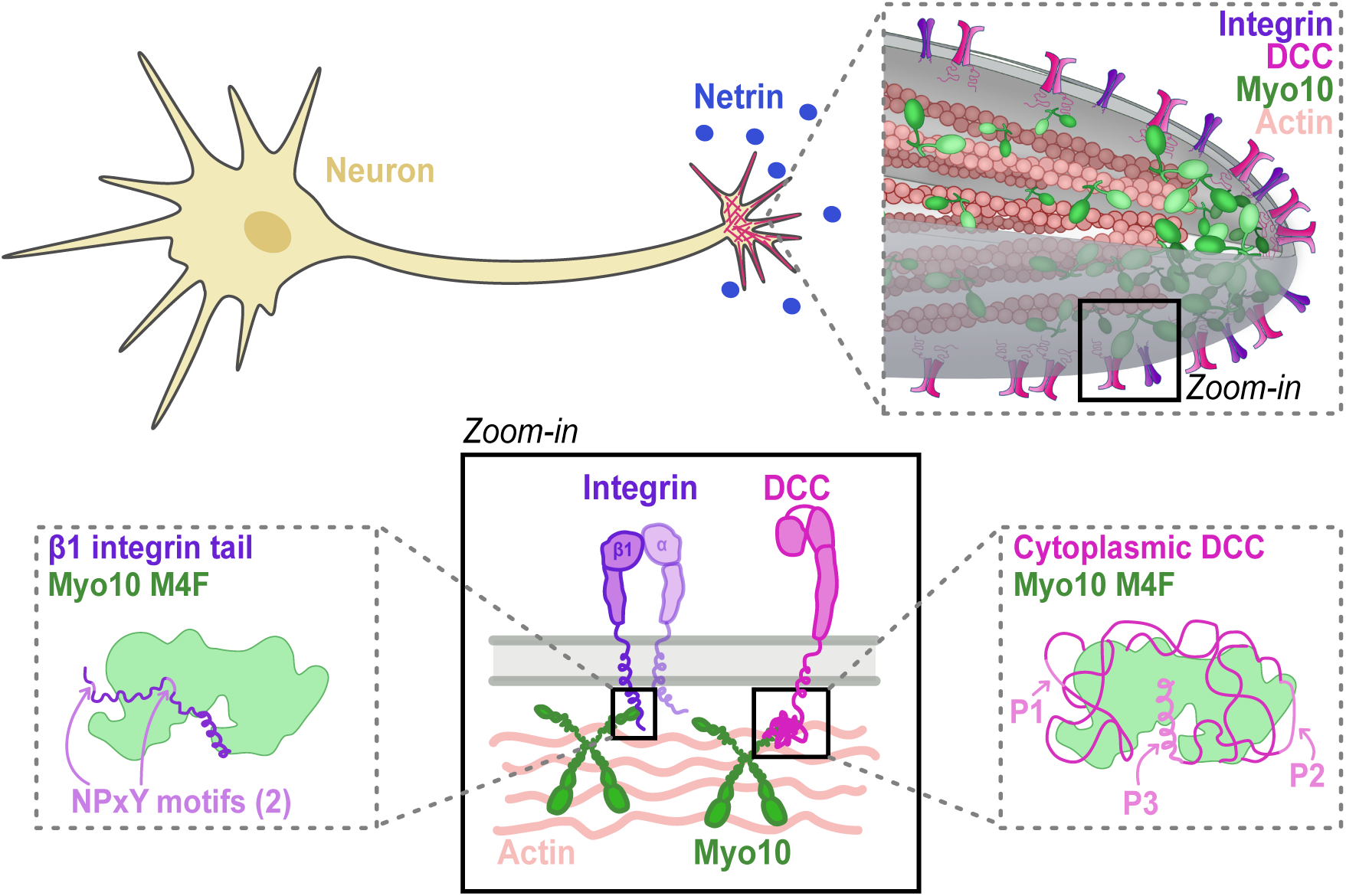
Myo10 interacts with cargos β1 integrin and DCC in filopodia. Myo10 promotes filopodia formation in cell types such as neurons, where axon guidance cues like netrin mediate directed axonal growth. At filopodial tips, Myo10 interacts with the protein receptors DCC and β1 integrin. The cargo-binding M4F domain of Myo10 is known to interact with the NPxY motifs of cytoplasmic β1 integrin and the P3 motif of cytoplasmic DCC.

In this study, we aimed to uncover the molecular mechanism of Myo10’s filopodial transport of β1 integrin and DCC, and the contribution of IDRs in this process. We conducted a detailed investigation into how Myo10 interacts with β1 integrin and DCC through its M4F domain. Using a combination of hydrogen-deuterium exchange mass-spectrometry (HDX-MS) and crosslinking MS (XL-MS), we mapped the binding interfaces of β1 integrin and DCC with the M4F. DCC’s P3 motif undergoes a disorder-to-order transition upon binding to Myo10. However, some DCC IDRs remain disordered yet still contribute to binding. This binding mode both enhances filopodial localization and influences the timing of DCC recruitment to nascent filopodia. Myo10’s M4F domain binds the cytoplasmic β1 integrin tail at both shared and distinct sites with DCC. Using super-resolution imaging, we find that Myo10 often exists as an apparent 2:1 complex with DCC, as expected if dimers of DCC are unresolved, but is more frequently dissociated from β1 integrin. The β1 integrin accumulates on the side of the filopodium nearest to the glass surface. Moreover, when both cargoes are present, DCC can outcompete β1 integrin and disrupt its filopodial tip localization. By studying Myo10’s interactions with cell-surface receptors such as DCC and integrin, we gain insight into the structural features of Myo10 and its cargo that underlie key processes in neuronal development and tumor progression. More broadly, our results highlight the functional versatility of IDRs in binding interactions, serving as extendable “bungee cords” that can absorb mechanical forces and hold protein complexes together even under fluctuating forces.

## Results

### DCC P3 is stabilized upon binding near C-terminus of Myo10 M4F

We first probed the native state dynamics of the DCC cytoplasmic domain and M4F using HDX-MS. HDX-MS is a nonperturbative isotope labeling method that reports on protein folding and the backbone stability of non-proline residues.^33–35^ Proteins can occupy higher energy states where the hydrogen bonds formed by the backbone are transiently broken due to thermal fluctuations. If the protein is transferred into a deuterium solution, some of the amide protons can exchange with deuterons before the hydrogen bond reforms. If a region of the protein has higher stability, its hydrogen bonds break and exchange less often than a site that is rarely hydrogen bonded. After deuterium labeling is stopped (i.e., quenched), the protein is digested into peptides. Because quenching occurs prior to cleavage, peptides retain the labeling pattern from the intact protein, which can be determined with liquid chromatography-MS (LC-MS). While we often refer to the exchange level for different peptides, it should be appreciated that these peptides originated from a native protein where the deuterium exchange occurred.

HDX experiments for cytoplasmic DCC were performed at pD 5 on ice (Supplementary Figure 1, 2), and for M4F at pD 7, 25 °C (Supplementary Figure 3A-B, D). For the M4F-DCC bound complex, experiments were conducted at pD 5 on ice (DCC peptides) and at pD 7 and 25 °C (more stable M4F protein) to maximize the dynamic range of the HDX measurements and capture both stable and unstable protein regions. HDX results were mapped onto AlphaFold-Multimer structures (PAE plot in Supplementary Figure 3).

We find that most of the cytoplasmic DCC lacks stable hydrogen-bonded structure. Most regions exchange with a rate matching the so-called “chemical” rate of an unstructured protein, by which k_chem_/k_obs_ = protection factor (PF) ∼1, where *k*_obs_ is the observed rate. Interestingly, peptides covering the P3 motif (DCC residues 1426-1447) exhibit a PF∼3 and ∼1 at the N- and C-terminus, respectively, suggesting that only the N-terminus forms helical structure and does so only 2/3’s of the time (Figure 2A, Supplementary Figure 4). In contrast, M4F is mostly composed of stable helices although of variable stability across its length. Some parts of the protein exchange extremely slowly with a protection factor >10^5^, which translates to a ΔG_HX_ >7 kcal/mol. Linker regions, such as covered by peptide 1545-1554, have little protection for the entire peptide (PF≤10, Supplementary Figure 6).

**Figure 2.**
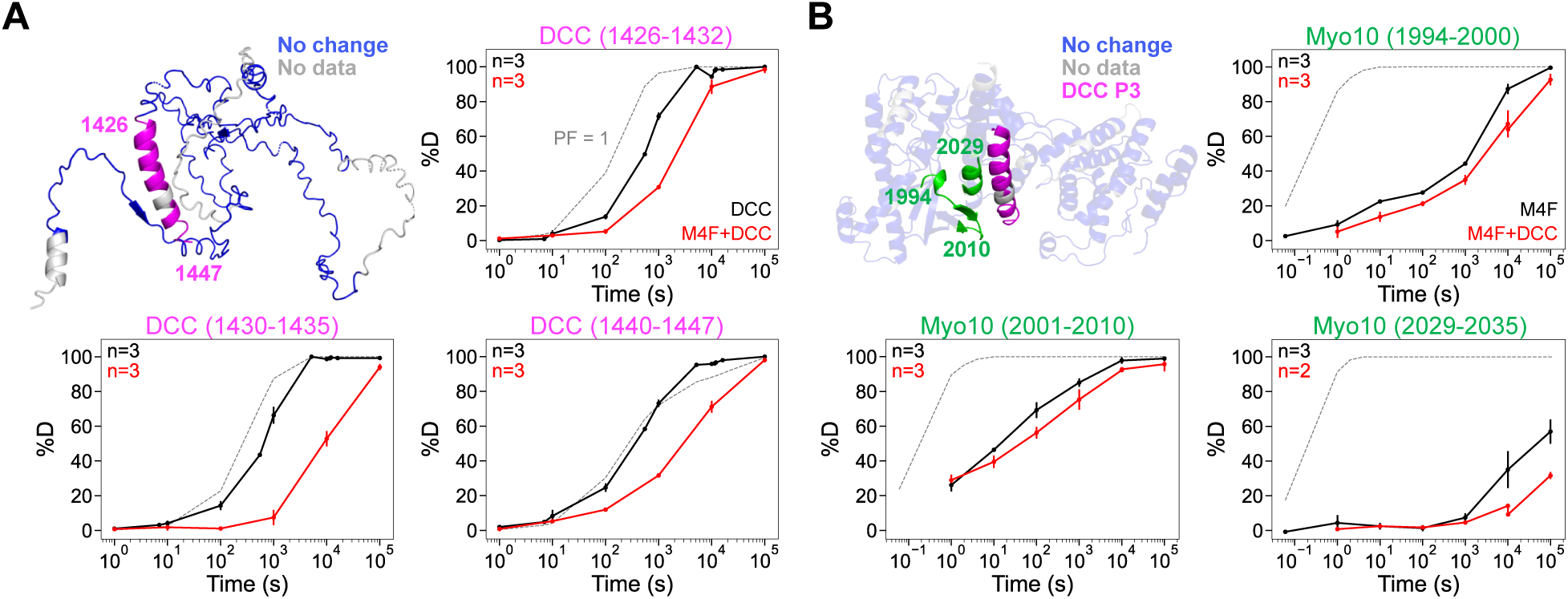
The DCC P3 helix and Myo10 M4F interface are stabilized upon binding. A) Deuterium uptake mapped on an AlphaFold-Multimer cytoplasmic DCC structure. Numbers next to the protein name indicate the residue positions on the full-length DCC sequence. Most of DCC is unchanged when bound to M4F (blue), except for the P3 helix (magenta) which gains 10x protection. The deuterium uptake plots represent coverage of the P3 helix (unbound = black line, bound = red line, k_chem_ = gray dashed line). B) Deuterium uptake mapped on an AlphaFold-Multimer M4F and cytoplasmic DCC structure. Numbers next to the protein name indicate the residue positions on the full-length Myo10 sequence. Most of the M4F is unprotected when bound to DCC (blue), except for peptides indicated in green that gain slight protection. The deuterium uptake plots represent coverage of the protected M4F peptides (as shown in (A)). Top left corner states the number of bioreplicates per condition. Data were normalized using in-exchange (0% D) and fully deuterated (100% D) controls; control data points are not shown. Points are means ±SD.

Next, we assessed the effects of cytoplasmic DCC and M4F binding on their stability and structure by HDX-MS. We expressed M4F and cytoplasmic DCC as a single-chain construct to increase their effective local concentration (C_eff_) and the fraction bound, thereby avoiding the need for high protein concentrations necessary to overcome their weak K_D_, while minimizing spectral crowding and improving signal-to-noise of MS data.^36^

The single-chain construct exhibits the same sites of M4F protection as HDX data of untethered cytoplasmic DCC and M4F, though with slightly different degrees of protection for M4F (∼5-30x higher PF compared to unbound, Supplementary Figure 5A). In the untethered form, there is no detectable protection anywhere for cytoplasmic DCC, further necessitating the use of the single-chain construct for the bound M4F-DCC measurements.

The P1 and P2 motifs of cytoplasmic DCC, along with most other regions, remain disordered and do not gain HDX protection upon binding to M4F (Supplementary Figure 4). However, about half of the residues in the DCC P3 motif have ∼10-fold greater protection (PF from ∼3 to ∼30), corresponding to a ΔΔ G_HX_ ∼1.4 kcal/mol (Figure 2A). HDX-MS performed on an M4F-DCCΔP1ΔP2 single-chain construct produces the same protection pattern for cytoplasmic M4F peptides, indicating that P1 and P2 do not influence the stability of the hydrogen bonds formed between M4F and the P3 helix (Supplementary Figure 5B).

Most of M4F exhibits no measurable change in PF upon cytoplasmic DCC binding except for a helix and β-strand where the PF increased by ∼3x at the site where DCC’s P3 helix docks (Myo10 peptide residues 1994-2000, 2001-2010, 2029-2035, as seen in structures of bound M4F-P3^11,15^) (Figure 2B, Supplementary Figure 6).

To examine the effect of electrostatic interactions between DCC and M4F, we performed HDX in low ionic conditions. DCC HDX experiments were originally conducted in 50 mM Na•acetate and 150 mM NaCl. We find that decreasing salt in the buffer to 10 mM Na•acetate and 2 mM NaCl does not noticeably change the HDX uptake curve between cytoplasmic DCC and M4F. This result indicates that electrostatic interactions are not major driving forces in their binding affinity (Supplementary Figure 5C).

### Disordered DCC regions also contribute to Myo10 binding

According to the HDX-MS results, only the P3 motif of cytoplasmic DCC increases its stability upon binding M4F, presumably in a helical conformation. Conversely, M4F gains stability proximal to the P3 helix (PF∼3). Since HDX-MS probes hydrogen bonding, we wondered if there could be non-hydrogen bond interactions between M4F and cytoplasmic DCC. Specifically, we wanted to determine whether disordered regions of cytoplasmic DCC still contribute to complex formation.

To capture weak and/or transient interactions, we performed crosslinking-MS (XL-MS) using a DSSO crosslinker that bridges lysines within 10 Å. DSSO crosslinking between M4F and DCC’s cytoplasmic domain reveals “hotspots” on DCC that react with multiple sites on M4F (Figure 3A, Supplementary Figure 7). Of note, one hotspot is in DCC’s disordered N-terminus (residue 1140, Supplementary Figure 8A), situated in a disordered protein binding domain predicted by DISOPRED3 (Supplementary Figure 8B), while another hotspot is in the P1 motif (residue 1167 of DCC). Other sites of crosslinking activity are in disordered regions downstream of the P2 motif, which has a PF∼1 (Supplementary Figure 7, residues 1380 and 1398 of DCC) (Figure 3A, Supplementary Figure 8A). Certain sites on M4F participate in more than one crosslink with DCC (Supplementary Figure 7), indicative of an ensemble of binding conformations. As seen in the AlphaFold-Multimer structure of the complex (PAE plot in Supplementary Figure 3), it would be impossible for these crosslinks to occur unless the DCC’s IDRs could transiently form alternative interfaces with M4F (Supplementary Figure 8C). Therefore, the XL-MS results highlight the flexibility of DCC’s IDRs to contact multiple sites of M4F.

**Figure 3.**
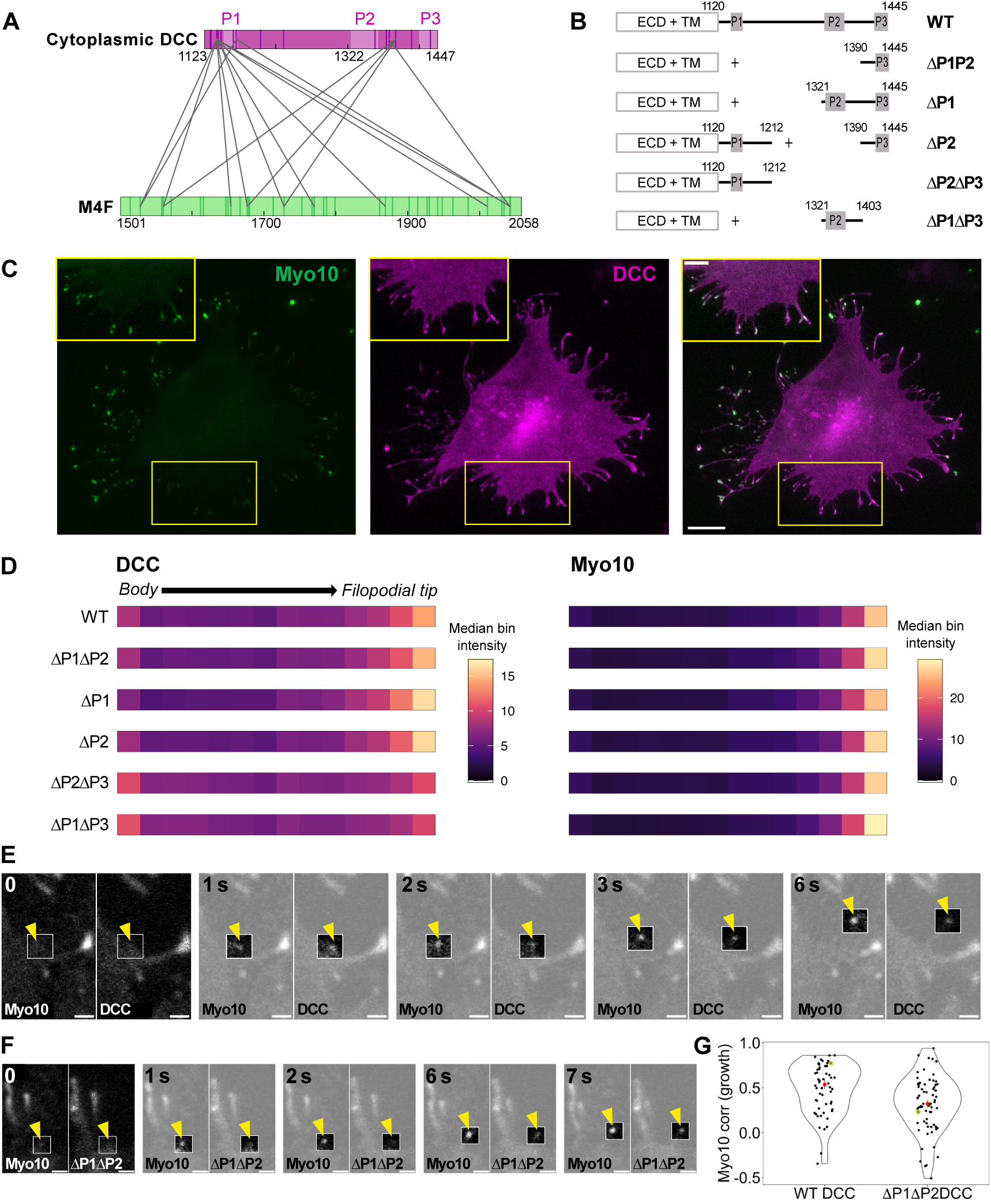
Disordered DCC regions contribute to Myo10 binding and affect association dynamics during filopodial initiation. A) Detected crosslinks between cytoplasmic DCC and M4F. Numbers correspond to the residue positions on the full-length DCC and Myo10 sequences. The vertical lines on each protein’s sequence represent lysines. Light pink regions annotate P1, P2, and P3 motifs. Note that DCC sites at the N-terminus, in P1, and downstream of P2 crosslink across M4F. B) DCC truncation mutations used in parts C-G. The entire full-length human DCC sequence was expressed with specific deletions in its cytoplasmic domain. C) Epifluorescence images of exogenously expressed HaloTag-Myo10 and EGFP-DCC in U2OS cells. Myo10 is labeled with HaloTag ligand-TMR (green) and DCC visualized by EGFP (magenta). Top right inset is a zoom-in of the yellow boxed cellular region. Both scale bars = 10 µm. D) Heatmap displaying the filopodial localization of DCC mutant constructs with the corresponding full-length Myo10 signal. Refer to methods for filopodial length and signal normalization procedure. Median values of each bin across n filopodia per 3 bioreplicates are displayed (WT, 1702 filopodia, 90 cells; ΔP2ΔP3, 1971 filopodia, 121 cells; ΔP1ΔP3, 1423 filopodia, 121 cells; ΔP1ΔP2, 652 filopodia, 93 cells; ΔP1, 1623 filopodia, 98 cells; ΔP2, 843 filopodia, 91 cells). Filopodial lengths > 2 µm were analyzed. E) Frame from a live movie of full-length DCC and Myo10. Yellow arrow points to a Myo10 and DCC co-localized punctum emerging from the cell membrane. Elapsed seconds from the first frame is depicted. Scale bar = 1 µm. F) Frames from a live movie of ΔP1ΔP2-DCC and Myo10. Yellow arrow points to a Myo10 punctum emerging from the cell membrane, where ΔP1ΔP2-DCC lags before accumulating with Myo10 at the filopodial tip. Elapsed seconds from the first frame is depicted. Scale bar = 1 µm. G) Distribution of Pearson correlation coefficients of intensity over time for Myo10 vs. DCC cargo. Each point represents one filopodium. Median ± SE (red) = 0.54 ± 0.05 (WT, 54 filopodia, 32 cells), 0.32 ± 0.04 (ΔP1ΔP2, 70 filopodia, 34 cells). Data are from 4 bioreplicates. Error bars are the SE for 5000 filopodial bootstrapped samples. Yellow circles are drawn around the events displayed in E and F.

Taken together, the HDX-MS and XL-MS experiments highlight that both intrinsic disorder, secondary structure formation (folding upon binding) and side-chain interactions can contribute to the affinity between DCC and Myo10. We propose that the flexibility of these IDRs allows DCC to remain associated with Myo10 when subject to impulses inside filopodia, illustrating how IDRs could act as elastic bungee cords.

### Disordered DCC regions affect association dynamics with Myo10 during filopodial initiation and elongation

To identify the minimal domains needed for their interaction in cells, we overexpressed full-length DCC and Myo10 in U2OS cells, along with DCC mutants lacking individual P motifs (Figure 3B). Netrin-1 was also expressed in the cells to provide a ligand for DCC activation and because of reports showing that netrin-1 prevents DCC-induced apopotosis.^37,38^ We mapped the localization of protein signal along filopodia. As expected, both wildtype (WT) DCC and Myo10 predominantly colocalize at the filopodial tip. Interestingly, the DCC P3 helix is sufficient for Myo10 binding, whereas the disordered P1 and P2 domains alone show weak but detectable interactions with Myo10 at filopodial tips (Figure 3D, ΔP1ΔP3 and ΔP2ΔP3). Deletion of any DCC P motifs does not produce dramatic changes in filopodial length (Supplementary Figure 8D).

Although HDX-MS indicates that the DCC P3 helix is the only region of cytoplasmic DCC with a measurable ΔΔG_HX_ upon M4F binding, we wanted to assess how the DCC P motifs contribute to filopodial processes in live cells using TIRF microscopy. WT DCC typically co-accumulates with Myo10 at the cell membrane and then travels together in newly formed filopodia (Figure 3E, Figure 3G). Unlike the WT, imaging the ΔP1ΔP2 DCC construct reveals a delay in co-accumulation with Myo10 during filopodial initiation (Figure 3F) and less correlated movement with Myo10 in growing filopodia compared to WT DCC (Figure 3G). This difference in co-accumulation suggests that the IDRs of DCC could recruit DCC to Myo10 clusters at membrane sites of new filopodia and help keep the complex associated at the onset of filopodial formation and during filopodial transport. The ΔP1ΔP2 DCC construct could be more prone to losing contact with Myo10 under mechanical force because it is structurally more brittle. In contrast, the bungee-cord like nature of P1, P2, and other cytoplasmic IDRs in WT DCC help maintain the integrity of the protein complex by allowing for extension of the motor-cargo linkage; if one interaction breaks during filopodial movement, another transient IDR-mediated interaction can form and keep the proteins connected. Therefore, the IDRs of DCC remain important by contributing multiple weak interactions and extensibility that collectively stabilize the complex.

### Myo10 M4F binds DCC and β1 integrin at both unique and shared sites

We pinpointed the peptides of M4F that undergo stabilization when DCC binds through HDX-MS and XL-MS (Figure 2, Figure 3A). It has been established that Myo10’s FERM domain binds both DCC and other cargos such as integrin.^20,39^ To compare M4F binding modes with different cargos, we performed HDX-MS on M4F with the cytoplasmic β1A tail, which is a common β1 splice variant (Supplementary Figure 3C & 3E, Supplementary Figure 9A-C). We performed β1 integrin tail HDX experiments at pD 5 on ice, and M4F-integrin bound complex experiments at pD 5 on ice (to analyze β1 integrin peptides) and at 25 °C (to analyze M4F peptides). The pD and temperature conditions were chosen to maximize the dynamic range of the HDX experiments. As done with bound DCC, we expressed M4F and β1 integrin tail as a single-chain construct to increase C_eff_ and the fraction bound. We inserted a short cytoplasmic DCC IDR sequence (40 residues) as a linker between the M4F and β1 integrin tail sequences; two versions of disordered DCC sequences yield similar results.

Interestingly, the membrane-proximal and membrane-distal NPxY motifs of β1 integrin tail, which are previously characterized motifs that bind Myo10, bind M4F in disordered conformations (Figure 4A, Supplementary Figure 10). Two regions on M4F exhibit slight protection upon binding the β1 integrin tail (increased ∼2x in PF vs. unbound M4F, Supplementary Figure 11). One contiguous region (containing Myo10 peptide residues 1994-2000 and 2001-2010) overlaps a region used by bound cytoplasmic DCC, while the second site is unique to the β1 integrin tail (Myo10 peptide residues 1557-1567) (Figure 4B). These peptides appear as potential binding sites in AlphaFold3 models of M4F bound to the β1 integrin tail (Supplementary Figure 9D). This suggests that Myo10 may generally dock cargo at one common site and then engage secondary sites to further stabilize specific cargo. On M4F, residues 1994-2000 correspond to a previously reported β1 integrin-binding region described by Miihkinen et al.,^14^ whereas residues 1557-1567 represent a newly identified interaction site. Our HDX-MS results broaden our understanding of how β1 integrin engages M4F.

**Figure 4.**
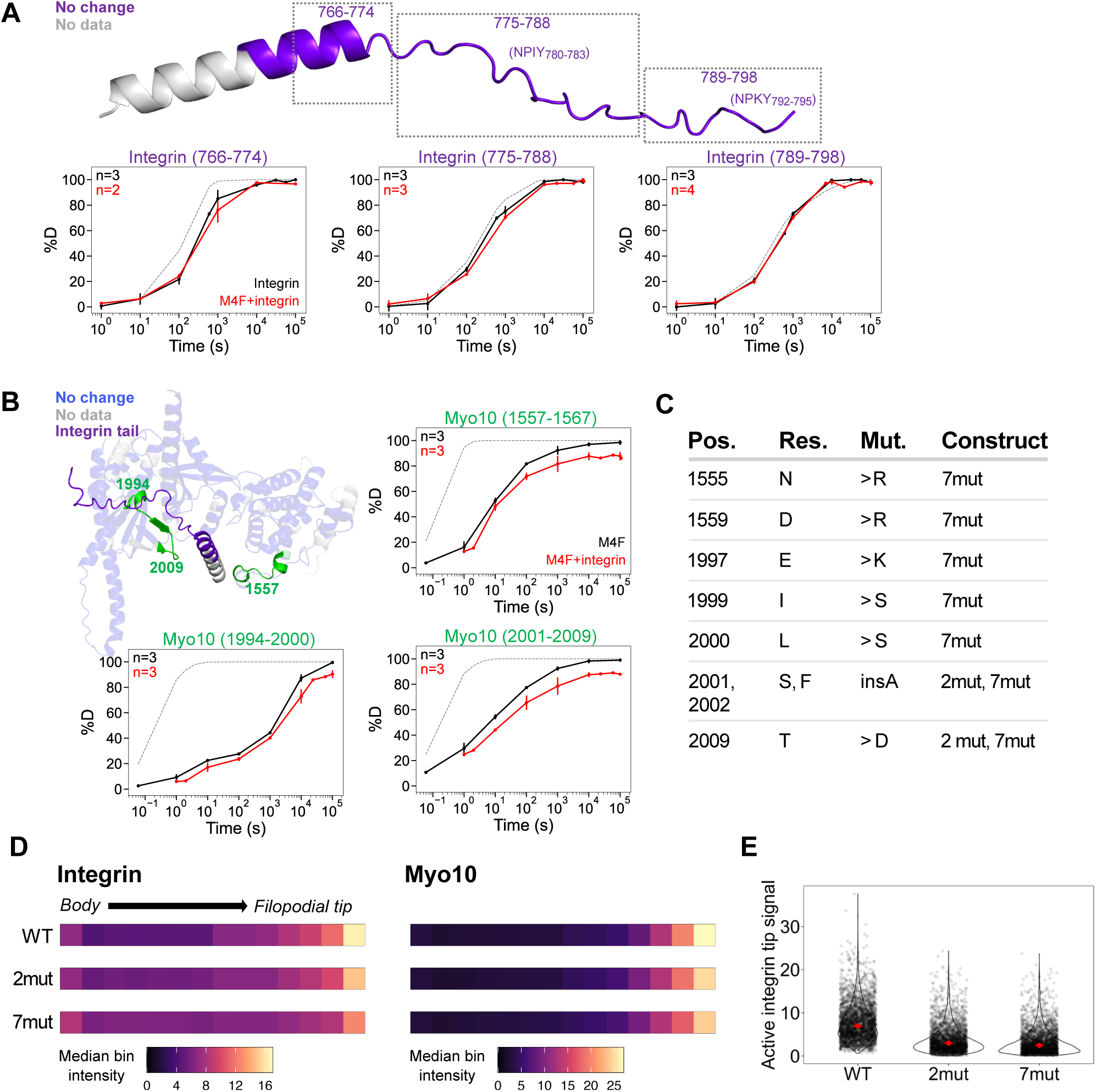
Myo10 M4F binds cytoplasmic β1 integrin at shared and unique sites compared to cytoplasmic DCC. A) Deuterium uptake mapped on an AlphaFold3 cytoplasmic β1 integrin structure. β1 integrin uptake is unchanged when bound to M4F. Numbers next to the protein name indicate the residue positions on the full-length β1 integrin sequence. The deuterium uptake plots represent example peptides across the sequence, revealing that the NPxY motifs remain unprotected during binding (unbound = black line, bound (single-chain construct) = red line, k_chem_ = gray dashed line). Top left corner states the number of bioreplicates per condition. Data were normalized using in-exchange (0% D) and fully deuterated (100% D) controls; control data points are not shown. Points are means ±SD. B) Deuterium uptake mapped on an AlphaFold3 M4F and cytoplasmic β1 integrin structure. Numbers next to the protein name indicate the residue positions on the full-length Myo10 sequence. Most of M4F is unchanged when bound to β1 integrin (purple), except for peptides indicated in green that gain slight protection. The deuterium uptake plots represent coverage of the protected M4F peptides (unbound = black line, bound (single-chain construct) = red line, k_chem_ = gray dashed line). Top left corner states the number of bioreplicates per condition. Data were normalized using in-exchange (0% D) and fully deuterated (100% D) controls; control data points are not shown. Points are means ±SD. C) Mutations in M4F intended to abolish binding between Myo10 and β1 integrin. Myo10-2mut comprises only SF2001-2002insA and T2009D, while Myo10-7mut contains all mutations in the table. D) Heatmap displaying the filopodial localization of Myo10 mutants and active β1 integrin. Refer to methods for filopodial length and signal normalization procedure. Median values of each bin across n filopodia per 3 bioreplicates are displayed (WT, 1443 filopodia, 115 cells; 2-mut, 493 filopodia, 42 cells; 7-mut, 687 filopodia, 45 cells). Filopodial lengths > 2 µm were analyzed. E) Signal intensity of active β1 integrin at the filopodial tip for WT Myo10, Myo10-2mut, and Myo10-7mut shown in part D. Medians ± SE (red) = 6.96 ± 0.10 (WT), 2.99 ± 0.05 (2mut), 2.46 ± 0.04 (7mut). Error bars are the SE for 5000 bootstrapped samples. One-way ANOVA shows significance (F(2, 6257.9) = 1292.7, p < 2.2×10⁻¹⁶). A post-hoc Games Howell test showed that Myo10-7mut had significantly lower normalized signal than Myo10-2mut (estimate = 0.57, 95% CI [0.39, 0.76], p = 4.3×10⁻¹²), and lower than WT (estimate = 4.41, 95% CI [4.20, 4.63], p < 2.2×10⁻¹⁶). Myo10-2-mut also showed significantly lower normalized signal than WT (estimate = 3.84, 95% CI [3.62, 4.06], p = 2.76×10⁻¹²).

Of note, the shape and time-scale of the HDX uptake curves of bound M4F peptide 1557-1567 and peptide 2001-2009 are very similar (Supplementary Figure 9E). When physically connected or adjacent residues have the same uptake curve, these regions have the same stability and likely exchange via the same unfolding event. Hence, we hypothesize that peptides covering residues 1557-1567 and 2001-2009 participate in the same cooperative binding conformation near the N-terminus of the β1 integrin tail, whereas peptide 1994-2000 likely independently interacts with the disordered C-terminal linker.

To determine whether the potential β1 integrin binding sites on M4F identified by HDX affect regulation of β1 integrin activity at filopodial tips, we overexpressed full-length Myo10 mutants in U2OS cells and stained for active β1 integrin (Figure 4C, D); U2OS contains moderate levels of endogenous β1 integrin. Two Myo10 mutants were tested, one containing the two mutations (*SF2001-2002insA, T2009D*) previously described by Miihkinen et al. (referred to as the *Myo10-2Mut*) and another (referred to as *Myo10-7Mut*) containing additional substitutions *(N1555R, D1559R, E1997K, I1999S, L2000S, SF2001-2002insA, T2009D*). To identify residues within M4F peptide residues 1557-1567 for mutation that would disrupt binding, we used two complementary approaches. First, given the structural similarity between the FERM domains of talin and M4F, we looked at interacting residues in AlphaFold3-predicted talin-integrin complexes, aligned the talin and M4F sequences, and then selected homologous residues in M4F for mutation. Second, we compared M4F sequence alignments across different myosins to identify non-conserved residues in Myo10 for mutation. In cells, active β1 integrin primarily localizes to filopodial tips when co-expressed with WT Myo10 (Figure 4D-E, Supplementary Figure 12A). This localization is significantly reduced with the *Myo10-2Mut* and further reduced with the *Myo10-7Mut* (Figure 4D-E), with no drastic impact on filopodial lengths (Supplementary Figure 12B).

### Myo10 frequently pairs with DCC in filopodia, while β1 integrin dissociates and deposits on the basal filopodial surface

We sought to understand in detail how Myo10 organizes relative to its cargo within filopodia. To do that, we used DNA-PAINT to spatially resolve Myo10 motor domains and cargo in U2OS cells with an average localization precision of ∼5 nm (by nearest neighbor analysis, i.e., NeNA^40^) (Figure 5A, B); we used HDBSCAN^41^ to cluster raw PAINT-localizations and approximated each cluster center as the position of a single protein molecule. We focused our analysis on central segments of the filopodium, where myosins and cargo are less crowded than at the tip and where active transport could occur. In Figure 5, we converted protein localizations to a cylindrical coordinate system for each filopodium to facilitate analysis. As expected, cargo is localized to the outer circumference of filopodia while Myo10 is localized to the interior (Figure 5C, D, Supplementary Figure 13A, B). Filopodia are flattened on average, with an ellipsoidal profile and peaks in the density of azimuth values (θ) at 0 and π (Figure 5C). Interestingly, we observe that Myo10 arranges with its cargo differently along the circumference and length of filopodia. DCC appears uniformly around the filopodium (Figure 5A, C), whereas β1 integrin is biased to the basal filopodia surface, closer to the cover glass (Figure 5B, D). Irrespective of cargo type, there is dense localization of Myo10 and cargo at the filopodial tip.

**Figure 5.**
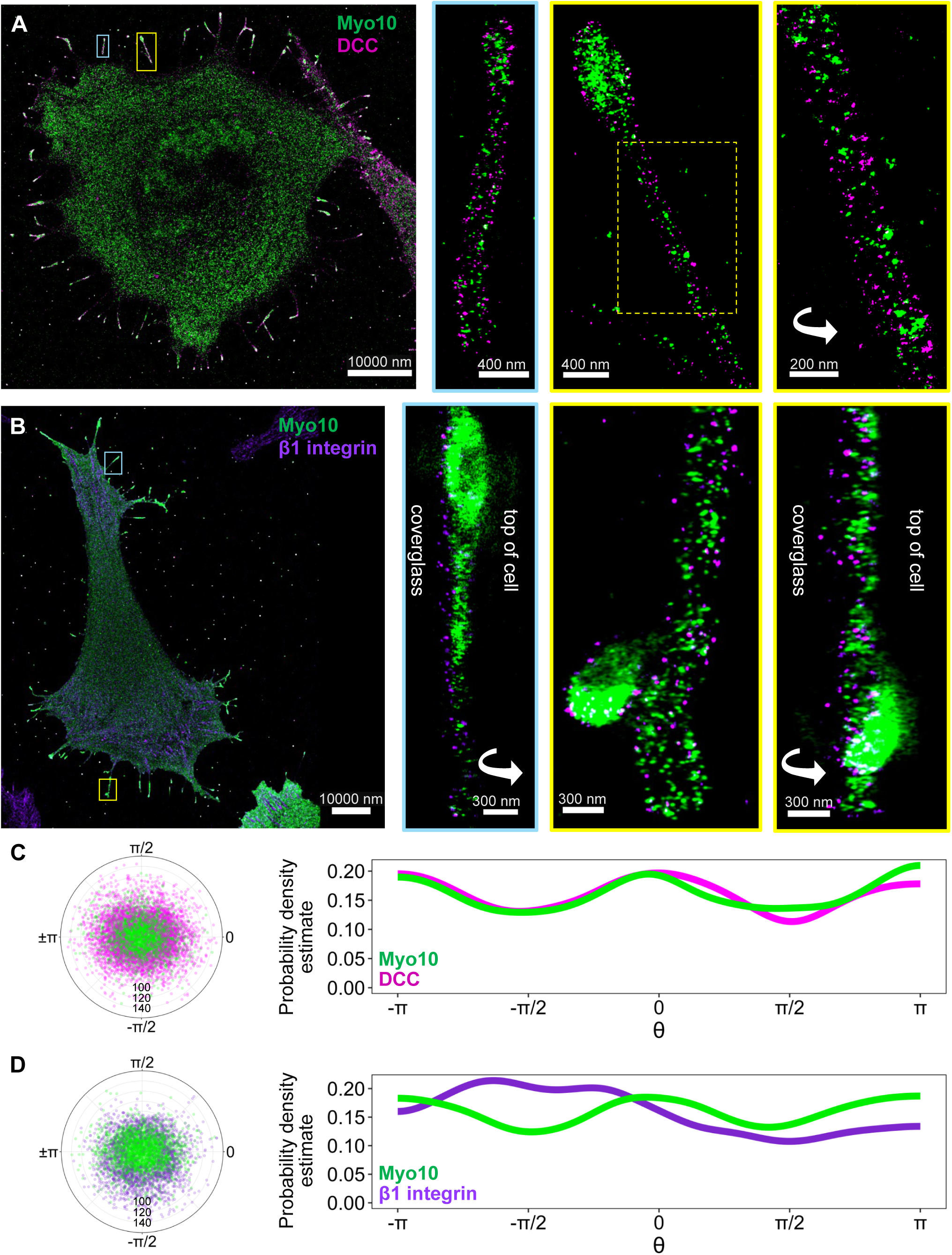
DCC organizes with Myo10 around the filopodial shaft, whereas β1 integrin is enriched at the basal filopodial surface. A) DNA-PAINT images of exogenously expressed HaloTag-Myo10 and EGFP-DCC in U2OS cells. Myo10 (green) was labeled with a HaloTag ligand-DNA conjugate and DCC (magenta) visualized by a nanobody-DNA conjugate directed at EGFP. Enlarged, representative filopodia (cyan, yellow boxes) are shown on the right. Scale bar = 10 µm (main). Rightmost image is a rotated view of the filopodium in the yellow box. B) DNA-PAINT images of exogenously expressed HaloTag-Myo10 and endogenous β1 integrin in U2OS cells. Myo10 (green) was labeled with a HaloTag ligand-DNA conjugate and β1 integrin (purple) visualized by a nanobody-DNA conjugate directed at 4B7R pan-β1 integrin antibody (purple). Enlargements of example filopodia (cyan, yellow boxes) are shown on the right. Scale bar = 10 µm (main). Zoom-in of the filopodium in the cyan box is rotated relative to its orientation in the original image. Rightmost image is a rotated view of the filopodium in the yellow box. C) An end-on view of filopodial protein localizations. Polar coordinate mapping of Myo10 and DCC molecule positions in 147 filopodial sections analyzed from 10 cells and D) Myo10 and β1 integrin molecules in 106 filopodial sections analyzed from 7 cells. Two-dimensional principal component analysis was performed, replacing the x and y coordinates with PC1 and PC2 while retaining the original z coordinate. Protein positions were projected onto the plane orthogonal to the filopodium’s principal axis (PC1) and converted to polar coordinates, with radius 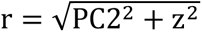 (black text labeling gray grid lines in nm) and azimuth 𝜃 = atan2(z, PC2) (in radians). Left: polar plot (filtered for r < 150 nm) of Myo10 and cargo clusters around the filopodial axis. Right: Distribution of 𝜃 of Myo10 and cargo in filopodia. Peaks at 0 and π reflect flattening of the filopodial cylinder. Note the enrichment of integrin from 0 to -π in (D).

Next, we investigated whether Myo10 and cargo concentrations were high enough to bind in the filopodial shaft (i.e., above their respective K_D_ values). To obtain protein concentrations, we estimated the volume of the analyzed filopodial region as a cylinder (in µm^3^), using the average of half the lengths of the sagittal and the transverse axes as the cylinder radius. Protein molecules per µm^3^ were then converted to µM. We corrected for the measured labeling efficiencies (LE) of 64% for the DCC probe (previously reported LE^42^) and 25% for Myo10 (Supplementary Figure 13C). We did not measure the LE of β1 integrin due to limitations in available probes for reliable quantification. Therefore, we report integrin concentrations calculated for 20% and 80% LE, reflecting typical bounds for DNA-PAINT probes^42^.

Along the length of the filopodia, Myo10 and DCC concentrations tend to be above their ∼0.5-2 µM K_D_,^11,15^ corresponding to a high fraction of bound DCC, whereas Myo10 and β1 integrin concentrations are more likely to be below their ∼25 µM K_D_^14^ with a lower fraction of bound β1 integrin (Figure 6A-E, Supplementary Figure 13D). Consistent with this picture, Myo10 and DCC appear to associate 2:1 (Figure 6A, B, F). We suspect that this apparent sub-stoichiometric association is due to challenges in resolving two nearby DCC molecules compared to two Myo10 motor domains that are separated on actin, and the actual Myo10:DCC ratio is 2:2. In contrast, the Myo10:integrin ratios vary considerably across filopodia, which is explained by Myo10 and β1 integrin being dissociated most of the time because their concentrations are below their K_D_. Alternatively, another interpretation is that Myo10 can carry multiple cargo molecules at a time.

**Figure 6.**
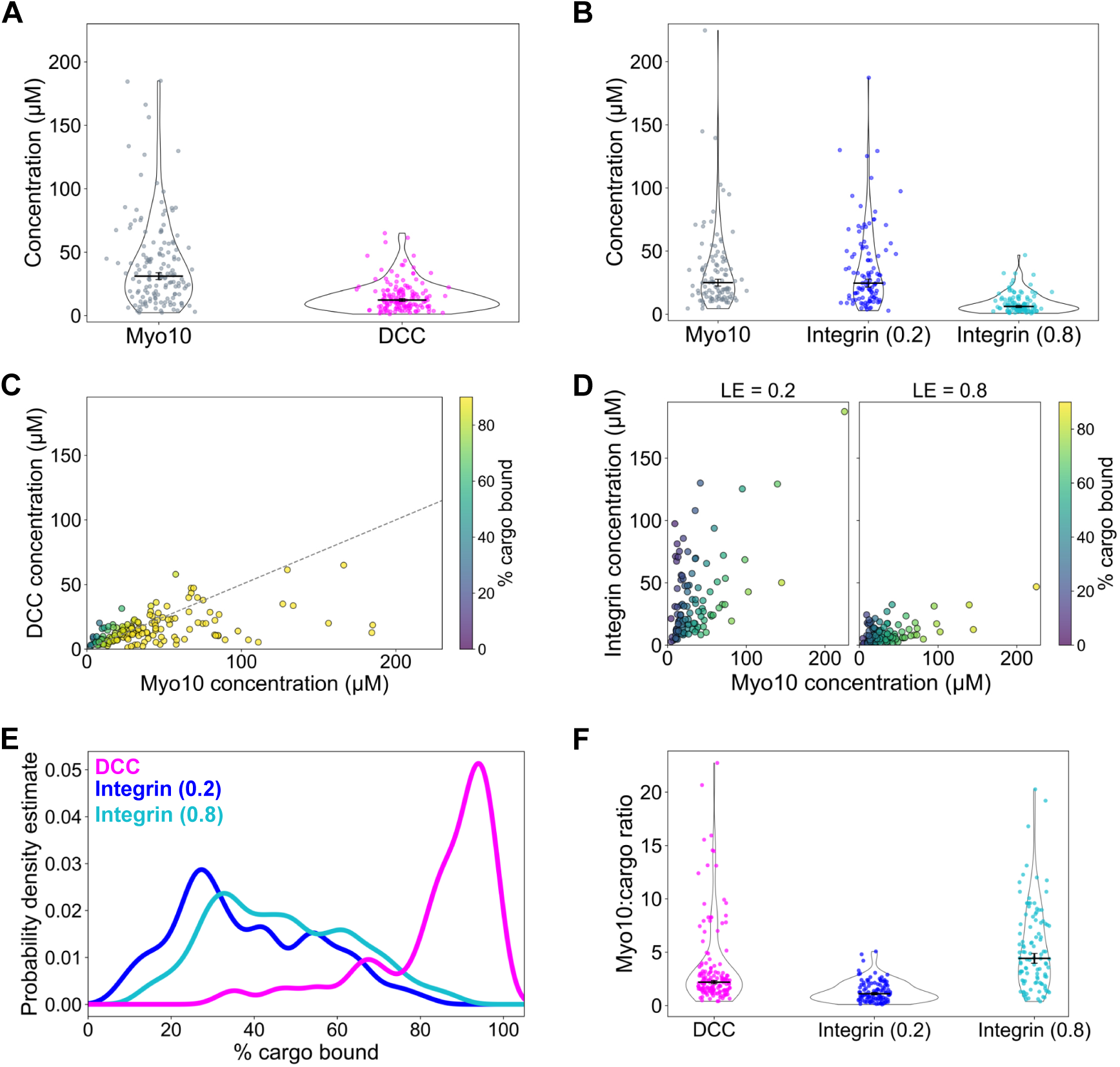
Cargoes are not fully occupied by Myo10 during filopodial transport. The local concentrations of Myo10 vs. cargo in the filopodial shaft are shown in A-F. To estimate the volume as a cylinder (µm^3^), the length of the longitudinal axis (i.e., longest side of the filopodial region) was used as the height, and the average of half the lengths of the sagittal and the transverse axis was used as the radius. Protein molecules per µm^3^ were converted to µM and corrected for labeling efficiency (LE) (HaloTag: 25%, DCC: 64%; Integrin: assumed efficiencies of 20% and 80% because unknown). See methods for details. All error bars are the SE for 5000 bootstrapped samples. A-B) Myo10 concentration distributions with DCC (magenta) or β1 integrin (dark blue, 20% LE; cyan, 80% LE) in filopodial shafts. Medians ± SE are indicated by horizontal black lines. Values: A) 31.13 ± 2.60 (Myo10, DCC dataset), 12.28 ± 1.10 (DCC). B) 24.98 ± 2.65 (Myo10, integrin dataset), 24.61 ± 3.11 (integrin at 20% LE), 6.15 ± 0.78 (integrin at 80% LE). C-D) Plots of Myo10 vs. cargo concentrations for DCC (C) and β1 integrin (D; left: 20% LE, right: 80% LE). Datapoints are colored by % of cargo bound by Myo10, calculated using the quadratic binding equation (K_D_: DCC = 2 µM; β1 integrin = 25 µM, published). Gray dashed line in (C) represents slope = 0.5. E) Distribution of % cargo bound by Myo10, calculated as in C-D. DCC in magenta; integrin in dark blue (20% LE) and cyan (80% LE). F) Ratio of Myo10 to cargo concentrations in filopodial shafts. Median ± SE (horizontal black lines) = 2.19 ± 0.14 (DCC), 1.10 ± 0.11 (integrin, 20% LE), 4.42 ± 0.45 (integrin, 80% LE).

To better understand the organization of cargo relative to Myo10, we attempted a mutual nearest neighbor analysis but were unable to generate fully convincing measurements. Mutual Myo10 and DCC nearest neighbors are usually closely associated (Supplementary Figure 13E). However, we obtain a similar distribution after randomization of the locations of one of the molecules in the filopodium (data not shown), so this measured distance largely reflects the density of molecules in the filopodial shaft rather than true interacting partners.

Additionally, we find little evidence of extension in the DCC IDRs. When considering that the stall force of Myo10 is ∼0.5 pN,^43^ we estimate a stretching of DCC IDRs by ∼13.5 nm using a wormlike-chain compliance of 0.038 pN/nm (Supplementary Figure 13F-G). Myo10 and β1 integrin are somewhat further apart, which is not surprising given that most integrins are most likely unbound (Supplementary Figure 13E).

### DCC competes with β1 integrin for Myo10 binding, resulting in repatterning of β1 integrin in filopodia

Integrin and DCC functions have been linked to certain cellular processes. Liu et. al. proposed that DCC could promote integrin signaling.^44^ Because β1 integrin and DCC both bind Myo10 in a similar location but with a different K_D_, we tested whether they could compete for Myo10 binding. To probe for potential binding competition, we overexpressed WT DCC and Myo10 in U2OS cells, stained for endogenous β1 integrin and then assessed the correlation between cargo and Myo10 fluorescence intensity along the entire length of filopodia (Figure 7). While β1 integrin alone concentrates at filopodial tips, coexpression with DCC leads to its frequent decoration along the length of the filopodium (Figure 7A, B). Overall, DCC has a consistently high correlation with Myo10 along filopodia. Active β1 integrin alone also has a high but somewhat weaker correlation with Myo10. However, when coexpressed with DCC, the active β1 integrin varies widely in its correlation with Myo10 (Figure 7C). Thus, DCC can outcompete β1 integrin for Myo10 binding in filopodia.

**Figure 7.**
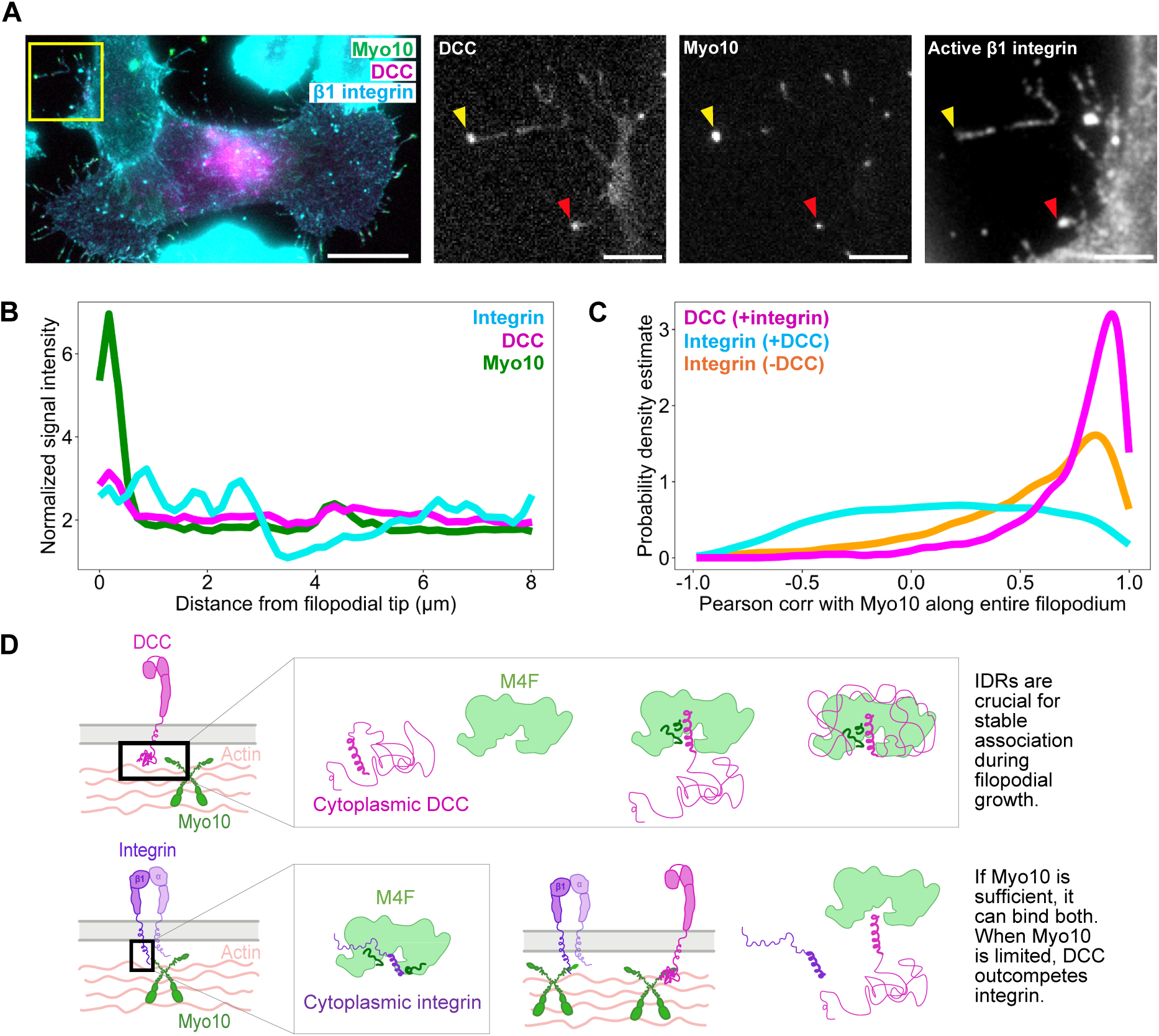
DCC competes with β1 integrin for binding to Myo10, causing a redistribution of β1 integrin in the filopodial shaft. A) Epifluorescence image of HaloTag-Myo10 (exogenous, green), EGFP-DCC (exogenous, magenta), and active β1 integrin (endogenous, cyan) in U2OS cells. Scale bar = 20 µm (main). Zoom-in of the yellow boxed region are shown for each protein channel. Yellow arrows highlight a punctum containing high Myo10 and DCC signal but low active β1 integrin signal (top arrow). Red arrows indicate a punctum containing high signal of all 3 proteins. Scale bar of enlargements = 5 µm. B) Signal intensity profiles of active β1 integrin, DCC and Myo10 of the filopodium annotated by the yellow arrow in part A. Each protein signal was normalized so that the sum of signal intensity across the filopodium = 100. C) Distribution of the Pearson correlation coefficient of a given filopodium’s Myo10 and cargo signal (DCC vs. active β1 integrin). Active β1 integrin (cyan) and DCC (magenta) correlations come from the same dataset, whereas active β1 integrin, without DCC present, (orange) comes from the dataset used in Figure 4D. Medians = 0.83 (DCC, +integrin), 0.15 (Integrin, +DCC), 0.64 (Integrin, -DCC). We analyzed n = 1946 (DCC+integrin), 1946 (Integrin+DCC), 2435 (Integrin-DCC) filopodia. D) Model of Myo10 binding to DCC compared to β1 integrin. Top: DCC P3 motif docks onto M4F (at dark green sites), bringing the rest of the disordered cytoplasmic sequence into proximity to wrap around M4F, holding the complex together during travel in filopodia. Bottom, left: β1 integrin tail docks onto M4F (at dark green sites), but this is a more brittle connection than Myo10 with DCC. Bottom, right: when DCC and β1 integrin are both present in filopodia, Myo10 can bind both cargos if in excess. Otherwise, Myo10 preferentially binds DCC.

## Discussion

Our study provides a compelling example of IDRs mediating protein binding. In the case of Myo10 and two types of cargo, we observe two distinct mechanisms by which IDRs and barely stable helices and turn motifs mediate interactions: disorder-to-order transition and “fuzzy” binding (i.e., no secondary structure when bound).^29^ According to the HDX-MS data, the P3 motif of DCC acts as a preformed recognition element whose conformational ensemble is shifted from being ∼30% unfolded to only ∼1% in the presence of Myo10. This aligns with previous results that describe a tunability of IDR sequences to encode weak secondary structures that influence binding kinetics and enable responsiveness to environmental changes.^45–47^

Besides the P3 motif, the other regions of cytoplasmic DCC maintain conformational heterogeneity and stay disordered upon binding M4F, exchanging with a PF∼1. The presence of alternative binding surfaces is evident from our XL-MS data, which reveal several contacts between DCC and M4F, especially at the N-terminus and near the P2 motif of DCC. Notably, the crosslinking hotspots at the N-terminus of cytoplasmic DCC make many of the same contacts with M4F as the hotspot downstream of the P2 motif. This observation could indicate the presence of two primary conformational states of DCC IDR binding to Myo10. An ensemble of bound configurations highlights the benefit of flexible IDR elements, allowing protein complexes to achieve high avidity through the sum of multiple weak interactions.

In our model (Figure 7D), the P3 helix brings DCC and M4F together with high specificity, serving as the most extensive point of contact between the two proteins; it is the only region that hydrogen bonds to M4F and is weakly stabilized upon binding according to HDX-MS. P1 and P2, along with other IDR motifs, contribute to the complex’s overall affinity and are likely important for establishing a stable interaction during the early stages of filopodial initiation, as shown by our live imaging results. We speculate that P1 and P2 could help with force-bearing because they can form numerous weak interactions that distribute mechanical load across the complex during cargo transport.

Whereas cytoplasmic DCC hydrogen bonds to M4F via a helical P3 motif, the β1 integrin tail binds via NPxY motifs that do not adopt stable secondary structure. We observe that two regions of M4F share the same HDX uptake curve shape and could cooperatively interact near the N-terminal half of the β1 integrin tail, while a third piece of M4F likely independently interacts with the C-terminal half of the β1 integrin tail. Appreciating that these systems are weak binding, we acknowledge that the observed increase in the PF of M4F is small (PF∼2) and could be near the limit of our experimental accuracy, but we note that this protection is not observed in other M4F peptides.

When we mutated the sites predicted to mediate Myo10 and β1 integrin binding, we observed less, though not completely abolished, integrin activation at the filopodial tip. Our *Myo10-7mut* only slightly decreases localization of active integrin at filopodial tips relative to the *Myo10-2mut.* We hypothesize that the three regions of M4F that dock the β1 integrin tail each exert a weak binding effect, and the overall interaction strength depends on the additive binding energies of these sites. Because the M4F and β1 integrin tail interaction is coordinated through multiple interactions, it is not surprising that removing an additional weak site only mildly reduces β1 integrin activation at filopodial tips. Furthermore, because integrin activation involves additional cofactors such as kindlin, the extent of reduction that we witness with the mutants suggests that other proteins make up for a binding-incompetent Myo10.

In addition to the regulation of integrin activity by talin and kindlin, mechanisms such as receptor clustering^48^ and applied force^49^ can regulate integrin activity. As such, Myo10 is only one of many pieces in the intricate network of integrin activation. The α-subunit of integrin could also compensate for reduced β1 integrin binding to Myo10.^14^ Integrin cytoplasmic tails serve as a landmark for multiple interaction partners, including SRC-family tyrosine kinases.^50^ Previous studies have described how integrin binds to talin via its membrane-proximal NPIY motif^51^ but talin is still sensitive to mutations at the membrane-distal NPXY motif.^18^ Furthermore, tyrosine-phosphorylation of β integrin tails modulate integrin’s affinity for talin as a means to regulate integrin activation.^51^ We cannot conclude from our study if Myo10 has particular affinities for an integrin NPxY motif or phosphorylation state. All these subtleties likely regulate Myo10 binding to integrin and determining their effects would be an enlightening future direction.

We also mapped specific regions where cargo docks to M4F. While we observe a general binding interface, short accessory peptides appear to reinforce specificity of the interaction. M4F shares some stabilized regions upon binding to either β1 integrin or DCC, but other sites are not the same, suggesting that Myo10 employs a tunable system of binding that allows selective recognition of distinct cargo, likely through side-chain mediated interactions that are not picked up by our HDX-MS experiments. A tunable binding system could explain how Myo10 can preferentially interact with a given target, depending on the cellular context and signaling needed.

From our super-resolution images, we observe a close association between DCC and Myo10 along and around the entire filopodial shaft, whereas β1 integrin appears to be ∼2x more likely to appear on the bottom of the filopodium at the cover glass. Due to technical limitations of our DNA-PAINT experiments, we cannot determine true Myo10-cargo interacting pairs (e.g., via nearest neighbor distances). The combined distances imposed by the placement of the protein labeling probes artificially expand the separation between Myo10 and cargo. As a result, the range of inferred binding distances includes the median nearest neighbor distances that we measured.

Based on protein concentration measurements, we hypothesize that Myo10 and DCC are frequently associated in an apparent 2:1 complex while traveling down filopodia. It makes sense that DCC and Myo10 are closely colocalized, considering that the cytoplasmic DCC IDR cannot be stretched far before reaching the 0.5 pN Myo10 stall force. It is possible that this low stall force is a specific adaptation of Myo10 for maintaining interactions between the DCC IDRs and M4F. In contrast to the association between DCC and Myo10, β1 integrin is frequently dissociated with a broad range of Myo10:integrin ratios. Note that this conclusion about the range of Myo10:integrin ratios is independent of any specific integrin labeling efficiency. Moreover, our conclusions of the bound fraction of integrin are robust to labeling efficiencies from 0.2-0.8. Unlike the cytoplasmic DCC IDRs, the β1 integrin tail is less compliant. This reduced compliance means that the M4F and β1 integrin tail interaction could be more prone to breaking during an impulse. This interpretation could explain our observation that integrin and Myo10 are less often associated along the filopodium, as integrin could be left behind and deposited along the bottom of the filopodia. On the basal surface, β1 integrin serves as an anchoring point for cellular adhesion. Alternatively, integrin and Myo10 might be less likely to associate along the filopodium because Myo10 primarily binds integrin at filopodial tips.

The observed range of Myo10:cargo association ratios, including the 2:1 ratio measured for DCC, support the proposal that Myo10 can associate with multiple cargo simultaneously, potentially through multivalent interactions that allow more efficient transport of protein oligomers. We previously observed Myo10 traveling in large clusters within filopodia,^52^ but the molecular nature of these higher-order assemblies remains unknown. If both Myo10 and its cargo can oligomerize, multivalency could contribute to avidity effects that substantially enhance the effective binding affinity beyond that predicted from one-site K_D_ models.^53^ Perhaps multivalency enables rapid cargo exchange to coordinate adhesion and guidance cues during cell migration or neurite extension.

Although most of our study focuses on Myo10 and a target cargo in isolation, we also witness an interesting interplay between β1 integrin, DCC and Myo10. DCC and integrin signaling pathways are connected through FAK. Xu et al. found that netrin-1-induced DCC clustering promotes P3 motif-mediated FAK dimerization, leading to PIP2-induced FAK clustering and FAK activation.^54^ Ren et al. observed that FAK colocalizes with DCC in rat cortical neurons, and perturbing FAK signaling disrupts netrin-1-induced neurite outgrowth and growth cone turning.^55^ Furthermore, Liu et al. suggests that DCC enhances integrin signaling and affects F-actin organization via integrin.^16^ DCC overexpression also promotes integrin-dependent FAK phosphorylation and basal F-actin organization. Meanwhile, talin is a known mediator of FAK activation downstream of integrin signaling,^56^ and FAK itself localizes near the plasma membrane within an integrin signaling layer in focal adhesions.^57^ Because we sometimes see high DCC signals coinciding with high active β1 integrin signal at filopodial tips, we do not rule out the possibility that the two proteins could promote each other’s activation, potentially through FAK. Given these connections, it would be valuable to further explore this connection between FAK, β1 integrin, and DCC in the context of Myo10-binding.

Scenarios in which both DCC and active β1 integrin concentrate at filopodial tips may also suggest that there is enough Myo10 to accommodate binding of all available cargo. At the filopodial tip, cargo competition for Myo10 binding is less of an issue because there is likely sufficiently high Myo10 concentration to bind all cargo. We do not know the precise number of molecules of cargo protein at the filopodial tips, but Myo10 can exist in large numbers (>500 molecules, median of ∼80 µM but can reach >500 μM).^52^ Therefore, this abundance of Myo10 could enable Myo10 to transport DCC and also activate integrins that have diffused to the same region. Our observation that active β1 integrin is more dispersed along the length of the filopodia when DCC is present aligns with integrin’s role in binding ECM, since integrin is likely responding to laminin on the coverslip. DCC appears to co-accumulate with Myo10 at specific membrane sites of newly budding filopodia prior to directed transport to filopodial tips by Myo10. Perhaps Myo10 deposits DCC at filopodial tips where DCC can interact with another ligand, netrin, before switching to bind β1 integrin that is present. Notably, the presence of DCC does affect active β1 integrin interactions with Myo10 (Figure 7C). We hypothesize that when there is cargo competition for Myo10 binding, Myo10 preferentially binds the cargo with stronger K_D_, which is DCC rather than β1 integrin. Although we did not perform DNA-PAINT imaging of cells co-expressing β1 integrin and DCC, we suspect that the β1 integrin is being deposited along the basal filopodial surface as seen in Figure 5 but at a much higher frequency when DCC is present. Regardless, the dynamics of Myo10’s cargo association are still not fully understood. Future studies could explore the mechanism of Myo10 on-and-off association with cargo and the equilibrium between swapping cargo partners.

How long-distance transport is achieved in thin cellular protrusions is still an open question. The most straightforward model is that a cytoskeletal motor picks up a cargo in the cell body and travels as a tightly-associated complex down the protrusion. Although the motor may occasionally detach and reattach to its actin track to start multiple processive transport runs, it does not release its cargo. In the other extreme, the motor transports itself without bound cargo to the end of the protrusion. There, it waits to capture cargo that diffuses into the protrusion. In this model, the motor protein has a patterning role rather than a transporting role. This patterning-capture process is possible for shorter protrusions such as filopodia (a few microns), but it is infeasible for long protrusions such as axons (up to meters).

Here, we find evidence for a third model, where transport occurs at concentrations near the K_D_ of the motor:cargo system. In this case, motors and cargoes can enter the protrusion but then dissociate and exchange while transport is happening. Each cargo would then undergo alternating periods of directed transport with free diffusion. Given the concentrations of each that we measure in filopodia, a Myo10:DCC complex is more likely than a Myo10:integrin complex. This preference of Myo10 for engaging DCC cargo is due to its higher affinity, but as we noted earlier, the longer DCC IDRs also play an important role in maintaining the connection. Although Myo10 is unlikely the main mechanism for integrin localization to filopodial tips, it plays a more significant role in DCC filopodial transport. Diffusion cannot be the only reason that DCC is highly localized in tips because the P3 deletion constructs show very weak tip localization, indicating that the DCC-Myo10 interaction is crucial in getting the bulk of DCC docked at the tip. Dynamic binding during transport might be important for patterning proteins along filopodia rather than concentrating everything at the filopodial tip, as we see with integrins and even more so with integrin when in competition with DCC.

A caveat of the DNA-PAINT images is that we are likely only looking at one type of “state” that filopodia are in because the analysis was conducted on regions of resolvable “single” molecule clusters without overcrowding. There are also filopodia that barely have any cargo or Myo10 in the filopodial shaft because they are all colocalized at the tip, which could be a result of Myo10 already having transported cargo there. Furthermore, we cannot conclude whether proteins in the filopodial shaft fell off a larger group during an earlier transport of a larger Myo10-cargo punctum. Because of the missing time component of the static images, we have no way of knowing whether Myo10 is heading out as part of the retrograde flow of actin or heading into the tip, or when cargo was deposited in the filopodial shaft. However, our live imaging of DCC and Myo10 provides initial insights into the dynamics of cargo binding. WT DCC puncta co-accumulate and co-move with Myo10 puncta from the membrane. Perhaps Myo10 typically transports “rafts” (i.e., DCC oligomers) out to tips. In this scenario, we would not be able to capture single DCC and Myo10 pairs and observe long processive runs with TIRF microscopy. However, that does not mean that Myo10 did not play a role in getting the cargo to the tip. Furthermore, we do not know the rate of “replenishing” or influx of single proteins into the filopodia. The filopodial transport system is complex, and future experiments will be needed to define these details.

Altogether, our findings reveal Myo10’s ability to decode IDR elements for coordination of diverse signaling cargos in filopodia. Our work suggests that tunable disorder within IDRs could be a general principle for cargo regulation in cellular transport. Testing our results in a physiological context of developing neurons could reveal how Myo10-mediated coordination of DCC and integrin signaling contributes to growth cone dynamics and neuron development.

## Acknowledgements

We thank Allen Huff for running the XL-MS samples and consulting on XL-MS analysis. This work was supported by the University of Chicago MCB Training Grant (T32 GM144292), University of Chicago Steinar Award 2023, and the NSF Graduate Research Fellowship (2140001) (to JS), IMPRS-ML (to SCMR), NIH grant and R01GM161048 (to RSR), and NIH R35 GM148233 (to TRS).

## Methods

### Plasmids and constructs

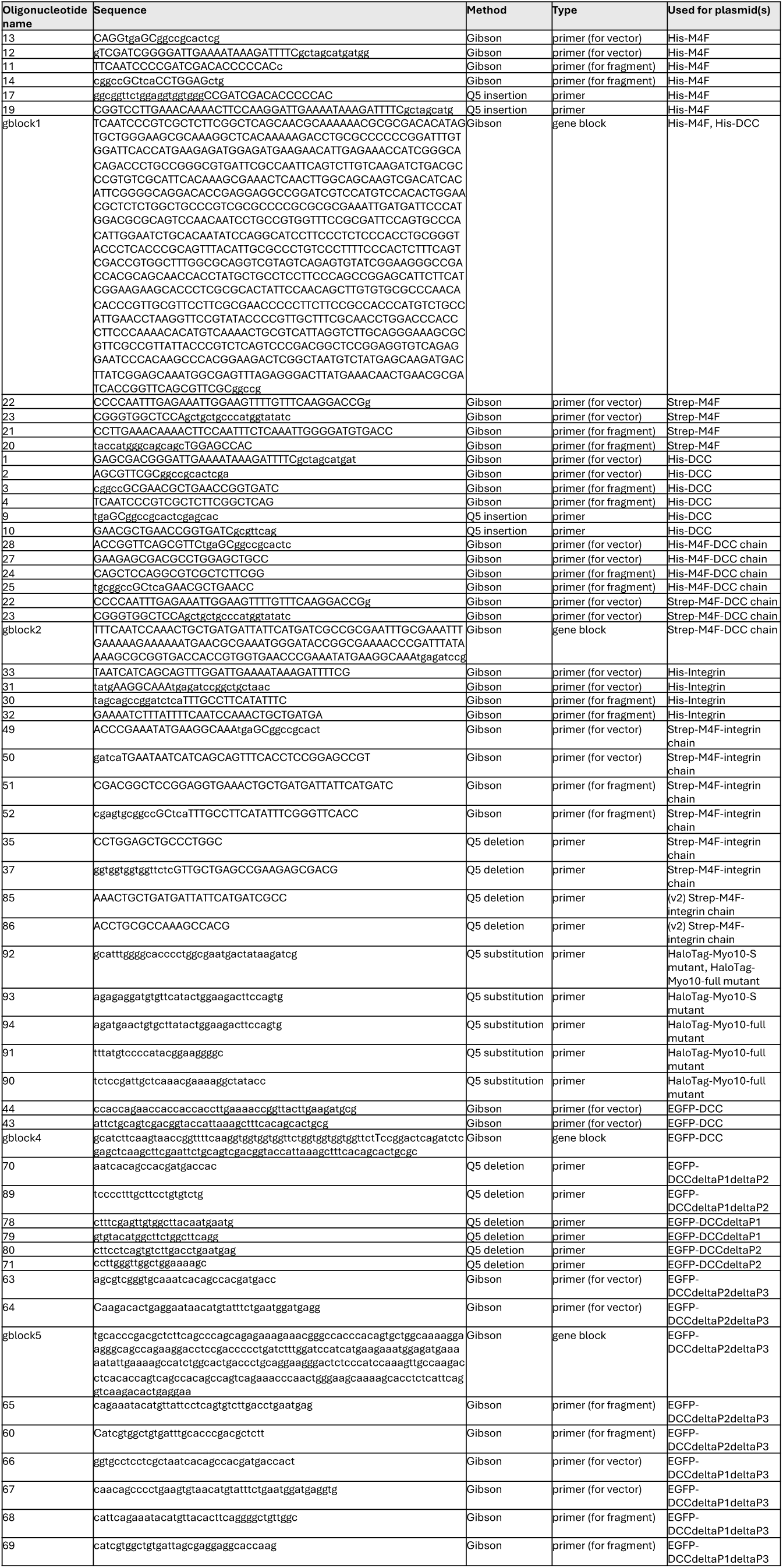

#### Constructs used for protein purification

His-M4F plasmid was generated in a pET28a backbone via Gibson Assembly and contains an N-terminal 8xHis-tag and TEV cleavage site. PCR products were cleaned and ligated via a KLD reaction. The pET28a backbone was amplified by PCR with primers 13 and 12 to produce the vector, and primers 11 and 14 were used to amplify the fragment containing cytoplasmic DCC sequence (gblock1). Then, a HRV3C cleavage site (TTGGAAGTTTTGTTTCAAGGACCG) and GS linker (ggcggttctggaggtggtggg) were inserted with primers 17 and 19 using the Q5 site-directed mutagenesis kit (New England Biolabs). PCR products were cleaned and ligated via a KLD reaction.

Strep-M4F plasmid was generated in a pET28a backbone via Gibson Assembly and contains an N-terminal 8xHis-tag and TEV cleavage site. The His-M4F plasmid was amplified by PCR with primers 22 and 23 to produce the vector, and primers 21 and 20 were used to generate the fragment containing beta1-integrin cytoplasmic tail sequence. Then, PCR products were assembled using Pfu Turbo mix.

His-DCC plasmid was generated in a pET28a backbone via Gibson Assembly and contains an N-terminal 8xHis-tag and TEV cleavage site. The pET28a backbone was amplified by PCR with primers 1 and 2 to produce the vector, and primers 3 and 4 were used to amplify the fragment containing cytoplasmic DCC sequence (IDT, gblock1). Then, PCR products were assembled using Pfu Turbo mix. A stop codon was added after the DCC sequence with primers 9 and 10 using a Q5 site-directed mutagenesis kit. PCR products were cleaned and ligated via a KLD reaction.

His-M4F-DCC chain plasmid was generated in a pET28a backbone via Gibson Assembly. The Strep-M4F plasmid was amplified by PCR with primers 28 and 27 to produce the vector, and primers 24 and 25 were used to amplify the fragment containing cytoplasmic DCC sequence (gblock1). Then, PCR products were assembled using Pfu Turbo mix.

Strep-M4F-DCC chain plasmid was generated via Gibson Assembly. The His-M4F-DCC chain plasmid was amplified by PCR with primers 22 and 23 to produce the vector and assembled with the fragment containing the TwinStrep tag and GS linker (IDT, gblock2). Then, PCR products were assembled using Pfu Turbo mix.

His-Integrin plasmid was generated in a pET21 backbone via Gibson Assembly and contains an N-terminal 8xHis-tag, Protein G (GATACCTATAAACTGGTGATTGTGCTGAACGGCACCACCTTTACCTATACCACCGAAGCGGTGGATGCGGCGACCGCGGAAAAAGTGTTTAAACAGTATGCGAACGATGCGGGCGTGGATGGCGAATGGACCTATGATGCGGCGACCAAAACCTTTACCGTGACCGAA), and TEV cleavage site. The pET21 backbone was amplified by PCR with primers 33 and 31) to produce the vector, and primers 30 and 32 were used to amplify the fragment containing beta1-integrin cytoplasmic tail sequence (gblock 2). Then, PCR products were assembled using Pfu Turbo mix.

Strep-M4F-integrin chain was generated in a pET28a backbone via Gibson Assembly. Strep-M4F plasmid (amplified with primers 49 and 50) served as the vector and then assembled with the β1-integrin cytoplasmic tail fragment (generated by His-integrin plasmid amplified with primers 51 and 52). Following sequence verification, primers 35 and 37 were used to shorten the linker between M4F and DCC via Q5 deletion, and the resulting PCR products were cleaned and ligated via KLD reaction.

To confirm that the M4F-integrin linker did not affect HDX-MS results, a second construct (Strep-M4F-integrin v2) was generated via two rounds of Q5 mutagenesis. The same Strep-M4F and His-integrin fragments (primers 49-52) were used, followed by mutagenesis with primers 85-86 and then primers 35-87, respectively. PCR products were cleaned and ligated using KLD reaction.

#### Constructs used for microscopy

HaloTag-Myo10-Flag plasmid was generated in a pTT5 vector backbone plasmid via Gibson Assembly and contains an N-terminal Halotag (Promega; GenBank: JF920304.1), full-length human Myo10 sequence (nucleotide sequence from GenBank: BC137168.1), and C-terminal Flag-tag (GATTATAAAGATGATGATGATAAA).

The HaloTag-Myo10-2mut was generated using the Q5 site-directed mutagenesis kit (New England Biolabs) to introduce the mutations of S2001 insA F2002 and 2009T to D (primers 92 and 93). PCR products were cleaned and ligated via a KLD reaction.

The HaloTag-Myo10-7mut was generated with two Q5 site-directed reactions. Primers 92 and 94 were used to introduce the mutations of 1997E to K, 1999I to S, 2000L to S, S2001 insA F2002, and 2009T to D. After sequencing confirmation, primers 91 and 90 were used to introduce the mutations of 1555N to R and1559D to R. PCR products were cleaned and ligated via a KLD reaction.

Netrin-Fc-His plasmid was acquired from Addgene (Addgene plasmid 72104).

EGFP-DCC plasmid was generated by modifying pCMV-DCC (Addgene plasmid 16459). The pCMV-DCC plasmid was amplified by PCR (primers 44, 43) and assembled with a gene block containing EGFP (IDT, gblock 4) via Gibson Assembly.

EGFP-DCCΔP1ΔP2 was generated using Q5 site-directed mutagenesis with primers 70 and 89 to delete residues 1120-1390 of the EGFP-DCC plasmid. PCR products were cleaned and ligated via a KLD reaction.

EGFP-DCCΔP1 was generated using Q5 site-directed mutagenesis with primers 78 and 79 to delete residues 1120-1321 of the EGFP-DCC plasmid. PCR products were cleaned and ligated via a KLD reaction.

EGFP-DCCΔP2 was generated using Q5 site-directed mutagenesis with primers 80 and 71 to delete residues 1212-1390 of the EGFP-DCC plasmid. PCR products were cleaned and ligated via a KLD reaction.

EGFP-DCCΔP2ΔP3 plasmid was generated via Gibson Assembly. The EGFP-DCC vector was amplified by PCR (primers 63, 64) and assembled with a gene block containing P1 (residues 1120-1212, IDT, gblock 5).

EGFP-DCCΔP2ΔP3 plasmid was generated via Gibson Assembly. The EGFP-DCC vector was amplified by PCR (primers 63, 64), and then assembled with a gene block containing P1 (residues 1120-1212, gblock 5) that was amplified by PCR (primers 65, 60).

EGFP-DCCΔP1ΔP3 plasmid was generated via Gibson Assembly. The EGFP-DCC vector was amplified by PCR (primers 66, 67) to produce the vector, and primers 68 and 69 were used to generate the fragment containing P2 (residues 1120-1212). Then, PCR products were assembled using Pfu Turbo mix.

All constructs were confirmed by Plasmidsaurus’s whole plasmid sequencing service using Oxford Nanopore Technology with custom analysis and annotation.

### Sample preparation for M4F

Three slightly different versions of M4F were used for 3 bioreplicates of M4F HDX-MS, all yielding similar results: His-M4F, M4F (His tag cleaved), and Strep-M4F.

Recombinant 8xHis-tagged M4F construct was overexpressed in E. coli strain BL21(DE3). Cells were lysed via sonication on ice, in lysis buffer (pH∼7.2) containing 50 mM HEPES, 200 mM NaCl, 5% glycerol, 1% igepal, 5 mM MgAcetate, 5 mM imidazole, 5 mM βME, 50 ug/mL DNase, 1 mg/ml lysozyme, 1 mM PMSF, and 1x protease cocktail inhibitor. Clarified lysate was loaded onto a buffer-equilibrated 5 mL Ni-NTA resin (Goldbio H-320-50) gravity-flow column at 4°C. Bound protein was first washed with 40 mL of buffer (pH∼7.7) containing 50 mM HEPES, 500 mM NaCl, 5% glycerol, 0.1% igepal, 25 mM imidazole, 5 mM βME, and 1 mM PMSF; then washed with 10 mL of buffer (pH∼7.7) containing 50 mM HEPES, 500 mM NaCl, 5% glycerol, 25 mM imidazole, and 5 mM βME; lastly, washed with 20 ml of buffer (pH=8) containing 50 mM HEPES, 500 mM NaCl, 5% glycerol, 50 mM imidazole, and 5 mM βME. Protein was eluted with 200 mM imidazole. Fractions containing the target protein were pooled and loaded into dialysis tubing for buffer exchange in 50 mM HEPES, 150 mM NaCl, 2 mM DTT, and 2.5% glycerol (pH∼8). HRV3C protease was added for removal of N-terminal His tag. After overnight dialysis at 4°C, sample was concentrated in a 30 kDa Pierce PES concentrator. For the M4F sample with His Tag intact, it was loaded onto a Superdex 200 10/300 GL size exclusion column for further purification.

Recombinant Strep-tagged M4F construct was overexpressed in E. coli strain BL21(DE3). Cells were lysed via sonication on ice, in lysis buffer (pH=8) containing 100 mM Tris, 200 mM NaCl, 1 mM EDTA, 5% glycerol, 1% igepal, 5 mM MgAcetate, 5 mM βME, 50 ug/mL DNase, 1 mg/ml lysozyme, 1 mM PMSF, and 1x protease cocktail inhibitor. Clarified lysate was loaded onto a buffer-equilibrated 3.5 mL Strep-Tactin resin gravity-flow column at 4°C. Bound protein was first washed with 25 mL of 50 mM Hepes, 500 mM NaCl, 0.1% igepal, 5% glycerol, and 1 mM PMSF, pH∼8; then 40 mL of buffer (pH=8) containing 100 mM Tris, 150 mM NaCl, 1 mM EDTA, 2.5% glycerol, 5 mM βME, and 1 mM PMSF; lastly, washed with 10 mL of buffer (pH=8) containing 100 mM Tris, 150 mM NaCl, 2.5% glycerol, 1 mM EDTA, and 5 mM βME. Protein was eluted with 50 mM biotin. Fractions containing the target protein were pooled and loaded into dialysis tubing for buffer exchange in 50 mM Tris, 125 mM NaCl, 1 mM EDTA, and 2 mM DTT (pH∼8). After overnight dialysis at 4°C, sample was concentrated in a 30 kDa Pierce PES concentrator.

### Sample preparation for DCC

Two bioreplicates of DCC (His tag cleaved) and one bioreplicate of 8xHis-tagged DCC were used for HDX-MS, all yielding similar results.

Recombinant 8xHis-tagged DCC construct was overexpressed in E. coli strain BL21(DE3). Cells were lysed via sonication on ice in the same lysis buffer used for the 8xHis-tagged M4F construct but with the addition of 6M GdmHCl. Clarified lysate was loaded onto a buffer-equilibrated 5 mL Ni-NTA resin gravity-flow column at 4°C and washed with the same buffers as described for 8xHis-tagged M4F construct purification, except that the first wash buffer contained 6M GdmHCl. Protein was eluted with 200 mM imidazole. Fractions containing the target protein were pooled and loaded into dialysis tubing for buffer exchange in 50 mM Tris, 125 mM NaCl, and 2 mM DTT (pH∼8). TEV protease was added for removal of N-terminal His tag. After overnight dialysis at 4°C, sample was concentrated in a 30 kDa Pierce PES concentrator. For one bioreplicate of DCC (His tag cleaved), it was loaded onto a Superdex 200 10/300 GL size exclusion column for further purification.

### Sample preparation for M4F-DCC chain

Two bioreplicates of Strep-M4F-DCC chain and one bioreplicate of M4F-DCC chain (His tag cleaved) were used for HDX-MS, all yielding similar results.

M4F-DCC chain was purified according to the His-tagged M4F construct methodology described earlier. After overnight dialysis and HRV3C cleavage at 4°C, sample was concentrated in a 50 kDa Pierce PES concentrator.

Strep-tagged M4F-DCC chain construct was purified according to Strep-M4F methodology described earlier.

### Sample preparation for integrin

Three bioreplicates of integrin (His tag cleaved) were used for HDX-MS.

Recombinant 8xHis-tagged integrin construct was purified similarly to the His-tagged M4F construct methodology described earlier. However, a 2.5 mL cobalt resin (TALON metal affinity, Takara 635501) resin gravity-flow column was used instead of nickel, and His-tag cleavage was done with TEV instead of HRV3C.

### Sample preparation for M4F-integrin chain

One linker version of Strep-M4F-integrin chain was used for 3 separate bioreplicates of HDX-MS, and a single bioreplicate of a slightly different linker version, all yielded similar results.

Strep-tagged M4F-integrin chain construct was purified similarly to the Strep-M4F methodology described earlier. Fractions containing the target protein were pooled and loaded into dialysis tubing for buffer exchange in 50 mM HEPES, 150 mM NaCl, 1 mM EDTA, 2.5% glycerol and 5 mM DTT (pH∼7.1). After overnight dialysis at 4°C, sample was concentrated in a 30 kDa Pierce PES concentrator. For one bioreplicate of Strep-tagged M4F-integrin chain (version 1 linker), it was loaded onto a Superdex 200 10/300 GL size exclusion column for further purification.

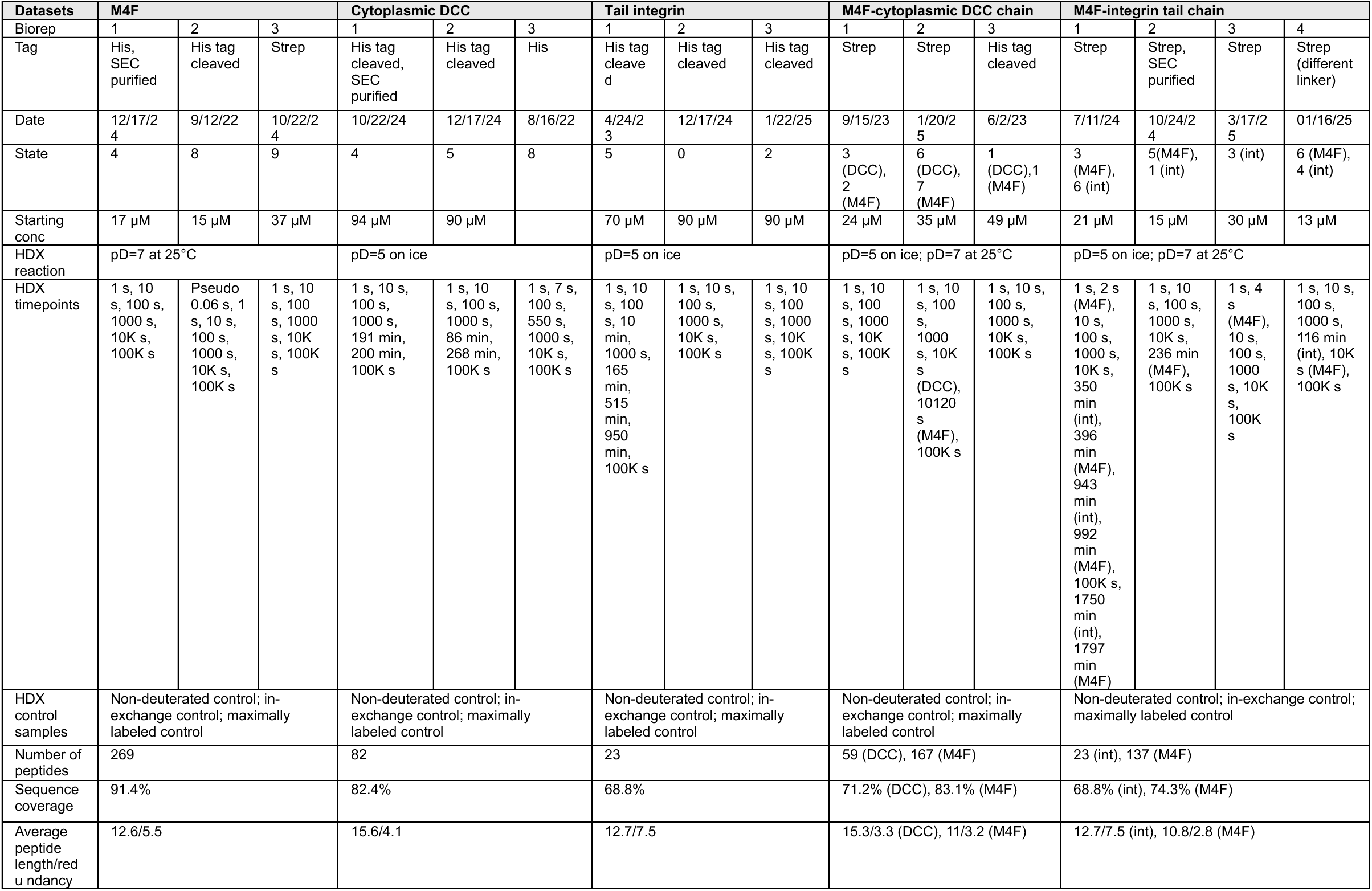

### Hydrogen-deuterium exchange labeling

HDX in a solution of ∼93% deuterium (D) content was initiated by diluting 2 μl of protein (starting concentration 15-80 μM in an H_2_O buffer) into 28 μl of D_2_O. Two HDX labeling conditions were used, depending on the protein target analyzed. For reactions conducted at pD=5 and 2°C, the deuterium buffer contained 50 mM NaAcetate and 150 mM NaCl; this was used for unbound DCC, M4F-DCC chain (for DCC peptide analysis), unbound integrin tail, and M4F-integrin chain (for integrin peptide analysis). For reactions conducted at pD=7 and 25°C, the deuterium buffer contained 30 mM HEPES and 150 mM NaCl; this was used for unbound M4F, M4F-DCC (for M4F peptide analysis), and M4F-integrin chain (for M4F peptide analysis). Reactions at pD=5 were performed in Eppendorf tubes sitting in a CoolRack on ice (2 °C). Timepoints included 1, 10, 100, 1000, 10k, and 100k seconds (with additional timepoints collected for some replicates, see table for details) before quenching with 30 μl of ice-chilled quench buffer (2 M urea, 600 mM glycine, pH=2.5) containing proteases.

In-solution digestion was optimized for each protein target. For unbound M4F, unbound DCC, M4F-DCC chain: 27 µL of quench buffer containing 2 µL of OD11 FPXIII and 1 µL of OD15 pepsin for a 3 min digestion on ice. For unbound integrin tail, M4F-integrin chain: 29 µL of quench buffer containing 1 µL of OD15 pepsin was incubated on ice for 2 min, followed by pipetting in 2.5 µL of OD11 FPXIII for another 2 min incubation (4 min in total). After digestion, samples were transferred to a 0.22 mm cellulose acetate filter column (Costar 8161) and spin-filtered at 15,000 RCF for 3 s at 2 °C prior to immediate injection into the LC-MS system. All HDX reactions were carried out in random order.

In-exchange controls (representing forward deuteration during the quenched reaction) were performed by mixing ice-chilled quench buffer and D_2_O buffer before addition to 2 µL of protein. Maximally labeled “full-D” controls were generated by incubating 2 µL of protein with pD=7 D_2_O buffer containing 2M urea (prepared with fully deuterated lyophilized urea powder) for ∼100k sec at 25°C.

MS/MS runs for peptide assignments and non-deuterated controls were carried out using the same protocol as above (with H_2_O buffers in replacement of D_2_O buffers).

### Hydrogen-deuterium exchange LC-MS set-up

After injection, peptides were desalted across a hand-packed trap column (Thermo Scientific POROS R2 reversed-phase resin 1112906, 1 mm ID × 2 cm, IDEX C-128) at 4°C for 160 sec, with a 100 μl/min flow of 0.1% formic acid (pH= 2.5).

Peptides were then separated on a C18 analytical column (TARGA, Higgins Analytical, TS-05M5-C183, 50 × 0.5 mm, 3 μm particle size) via a 13 min, 10–90% (vol/vol) acetonitrile (0.1% formic acid) gradient applied by a Dionex UltiMate-3000 pump. Eluted peptides were analyzed by a Thermo Q Exactive mass spectrometer (scan range of 400-2000 m/z, 140k resolution, and AGC target of 3e6).

### Hydrogen-deuterium exchange data analysis

For peptide assignment, MS/MS data was searched against sequences of target proteins, proteases, decoys, and other proteins run on our LC–MS system using SearchGUI version 4.0.25 (CompOmics Group) with the following settings: unspecific cleavage; precursor charge 1-8; isotopes 0-1; precursor m/z tolerance 10.0 ppm; fragment m/z tolerance 10.0 ppm; no post-translational modifications; peptide length 5-30. Search algorithms employed in SearchGUI included X! Tandem, MS Amanda, and Comet. Results were imported into PeptideShaker (version 2.0.18) (CompOmics Group) and later processed in EXMS2 to generate a peptide list. MS1 data was imported into HDExaminer 3.1 (Sierra Analytics) to fit peptide isotope distributions. Downstream plotting was done in a custom Jupyter Notebook.

### Cross-linking mass-spectrometry experiments

#### DSSO cross-linking reaction

Fifty micrograms of the M4F-cytoplasmic DCC complex (1:1 molar ratio) was prepared in 10 µL of reaction buffer (20 mM HEPES, 150 mM NaCl, 1.5 mM MgCl₂, 1 mM TCEP, pH 7.5) and equilibrated for 30 min at 25 °C. The solution was brought to 50 µL (1 mg/mL final) and incubated with an 80-fold molar excess of disuccinimidyl sulfoxide (DSSO; Cayman Chemical, 9002863; 100 mM stock in anhydrous DMSO). Cross-linking proceeded for 1 h at 25 °C and was quenched with 1 µL of 1 M Tris-HCl (pH 8.0). One biological replicate was analyzed in triplicate, and a single run was performed on another biological replicate. Cross-linked products were verified by SDS–PAGE under reducing conditions to confirm cross-linking efficiency.

#### Trypsin/Lys-C digestion and sample clean-up

Cross-linked proteins were reduced with 20 mM TCEP for 30 min at 65 °C and cooled to room temperature. Alkylation was performed with freshly prepared iodoacetamide (80 mM final) for 30 min in the dark. Samples were acidified with 5 µL of 12% phosphoric acid and diluted with 300 µL of S-Trap binding buffer (100 mM TEAB in 90% methanol, pH 7.55). Proteins were captured on S-Trap micro columns (Protifi) by centrifugation at 4000 × g for 30 s. The columns were washed three times with 200 µL of binding buffer. Digestion was performed by adding 20 µL of digestion buffer (50 mM TEAB, 0.5 mM CaCl₂, pH 8.0) containing Trypsin/Lys-C mix (1:10 enzyme-to-protein ratio, w/w) and incubating overnight at 37 °C in a humidified chamber. Peptides were sequentially eluted with 40 µL of (i) 50 mM TEAB, (ii) 0.15% formic acid in water, and (iii) 0.15% formic acid in 60% acetonitrile, each by centrifugation at 4000 × g for 1 min. The combined eluates were dried under vacuum and resuspended in 0.15% formic acid before LC-MS analysis.

#### LC-MS analysis

Peptide mixtures (400 ng) were analyzed on an UltiMate 3000 RSLCnano system (Thermo Fisher Scientific) coupled to an Orbitrap Eclipse Tribrid mass spectrometer and Nanospray Flex ion source. Peptides were loaded onto an Acclaim PepMap 100 C18 trap column (Thermo Scientific, 0.75mm ID × 2 cm, 3 µm particles) and separated on a 50 cm MonoCap column (GL Sciences, 0.75 mm in inner diameter; Cat. No. 5020-10006) was employed. The flow rate was maintained at 500 nL/min, and the temperature was held constant at 25 °C.

During the analysis, mobile phase A (0.15% formic acid in water) and mobile phase B (0.15% formic acid in 100% acetonitrile) were utilized. A gradient spanning 145 minutes was employed: 5% B for 5 minutes, followed by a transition from 5% to 20% B over 100 minutes, a transition from 20% to 32% B over 19 minutes, and a rapid transition from 32% to 95% B in 1 minute. The composition was maintained at 95% B for 5 minutes, then re-equilibrated at 4% B.

The mass spectrometer was operated in positive nanospray ionization mode at 2.0 kV with a static spray voltage, an ion-transfer-tube temperature of 305 °C, and RunStart Easy-IC internal calibration enabled. Data were acquired in data-dependent mode over a total 140-min run, with standard pressure and peptide application settings. Full MS scans were collected in the Orbitrap with wide quadrupole isolation across m/z 350–1500 at 60,000 resolution (at m/z 200), AGC target = 4 × 10⁵, maximum injection time = 50 ms, RF lens = 30 %, ensuring at least six points across each chromatographic peak. During each 5 s duty cycle, precursors with charge states 2–8 and intensities > 2 × 10⁴ were selected for fragmentation; dynamic exclusion was applied after one observation for 45 s within ±10 ppm, and isotopic peaks were excluded. For MS², precursor ions were isolated in the quadrupole (1.6 m/z window) and fragmented by collision-induced dissociation (CID) at fixed 25 % normalized collision energy (activation time = 10 ms, activation Q = 0.25). Resulting fragments were analyzed in the Orbitrap at 30 000 resolution with AGC target = 5 × 10⁴, maximum injection time = 54 ms, and profile mode. DSSO-specific cross-linked precursors were further subjected to MS³ acquisition triggered by a Δ = 31.9721 Da mass shift; re-isolated ions (1.6 m/z window, 3 m/z MS² isolation window) were fragmented by CID at 35 % energy (activation time = 10 ms, Q = 0.25) and detected in the ion trap at rapid scan rate (centroid mode, AGC target = 2 × 10⁴, maximum injection time = 120 ms). Parallel ETD MS² scans employed calibrated charge-dependent ETD parameters for precursors of charge 2–7 and were recorded in the Orbitrap at 30,000 resolution (m/z 150–2000, AGC target = 5 × 10⁴, maximum injection time = 100 ms). All spectra were acquired in positive polarity with source fragmentation disabled, ensuring comprehensive MS²–MS³ coverage of DSSO-cleavable cross-links while maintaining ≥ 6 data points per chromatographic peak.

Raw LC–MS/MS data were analyzed in Thermo Proteome Discoverer 3.0.1.27 using the *PWF_Fusion_Basic_SequestHT_XlinkCleavable_MS2_MS2_MS3* workflow followed by the *CWF_Basic_Xlinkx.pdConsensusWF_DSSO* consensus workflow. Two parallel Sequest HT searches were used. The CID-based search used Trypsin (full), up to two missed cleavages, peptide lengths of 6–150 aa, a precursor tolerance of 10 ppm, fragment tolerance of 0.02 Da, and ion weights of b = 1 and y = 1. The EThcD-based search used identical enzyme and mass tolerances but ion weights of b = 0.5, c = 1, y = 0.5, and z = 1. Dynamic modifications included Oxidation (+15.995 Da, M), DSSO-hydrolyzed (+176.014 Da, K), DSSO-Tris (+279.078 Da, K), and N-terminal Acetylation (+42.011 Da); Carbamidomethylation (+57.021 Da, C) was static. PSMs were validated using a target–decoy approach with 1% (strict) and 5% (relaxed) FDR and filtered to High confidence.

Crosslink identifications were performed using XlinkX with an MS2–MS2–MS3 acquisition strategy. DSSO (+158.004 Da on K) was specified as the crosslinker, with a minimum S/N of 1.5 and protein N-terminal linkage enabled. XlinkX Search used Trypsin (full), two missed cleavages, minimum peptide length of 5, a 10 ppm precursor tolerance, and FTMS/ITMS fragment tolerances of 20 ppm and 0.5 Da, respectively, with Carbamidomethyl (+57.021 Da, C) as static and Oxidation (+15.995 Da, M) as dynamic. Crosslinks were filtered and validated at 1% FDR, and the consensus workflow consolidated high-confidence identifications. High-confidence cross-links were defined by XlinkX Score > 100.

### Mammalian cell culture and transfection for microscopy

U2OS cells (ATCC HTB-96, tested negative for mycoplasma by PCR and DAPI staining) were passaged every 2-3 days and used under passage number 15 after thawing. Cells were grown at 37°C and 5% CO_2_ in Gibco 1x DMEM (Thermo Fisher 11995073) supplemented with 10% fetal bovine serum.

At 90-95% confluency, U2OS cells in one well of a 6-well dish were transiently transfected using either Lipofectamine 2000 or 3000 (see table for details). The well was replaced with 2 mL of pre-warmed DMEM + 10% FBS prior to transfection.

For Lipofectamine 2000, plasmid DNA was diluted in 250 µL Opti-MEM. In another tube, Lipofectamine 2000 was diluted at a 2.5:1 Lipofectamine:DNA ratio (6.75 µL Lipofectamine 2000 in 250 µL Opti-MEM), mixed, and incubated for 5 min. The two solutions were combined, mixed thoroughly, and incubated for 20 min at 25°C.

For Lipofectamine 3000, plasmid DNA was diluted in 125 µL Opti-MEM with P300 reagent at a 2:1 P300:DNA ratio. In another tube, Lipofectamine 3000 (2.5:1 Lipofectamine:DNA ratio) was prepared in 125 µL Opti-MEM, thoroughly vortexed, then combined with the DNA mixture and incubated 15 min at 25°C.

The resulting transfection complex (500 µL for Lipofectamine 2000; 250 µL for Lipofectamine 3000) was added dropwise to one confluent well of cells and gently rocked by hand to ensure even distribution. Cells were incubated at 37 °C with 5% CO₂. For Lipofectamine 2000 transfections, media was replaced with 2 mL DMEM + 10% FBS five hours post-transfection. Twenty-four hours after transfection, cells were seeded onto laminin-coated (20 µg/mL) 8-well chambered coverslips (for fixed cell samples) or 10 mm micro-well (round coverslip; for live cell samples) at 50%-60% density.

Three bioreplicates were conducted for each experiment, where transfection took place in separate wells and/or days.

| **Plasmids** | **Experiment** | **Transfection reagent** |
| --- | --- | --- |
| 2 µg of HaloTag-Myo10-FlagTag, 2 µg of EGFP-DCC, 1 µg of netrin | DNA-PAINT, Colocalization heatmap, Cargo correlation plots | Lipofectamine 2000 |
| 2 µg of HaloTag-Myo10-FlagTag, 2 µg of EGFP-DCCΔP1ΔP2, 1 µg of netrin | Colocalization heatmap | Lipofectamine 3000 |
| 2 µg of HaloTag-Myo10-FlagTag, 2 µg of EGFP-DCCΔP1, 1 µg of netrin | Colocalization heatmap | Lipofectamine 3000 |
| 2 µg of HaloTag-Myo10-FlagTag, 2 µg of EGFP-DCCΔP2, 1 µg of netrin | Colocalization heatmap | Lipofectamine 3000 |
| 2 µg of HaloTag-Myo10-FlagTag, 2 µg of EGFP-DCCΔP2ΔP3, 1 µg of netrin | Colocalization heatmap | Lipofectamine 2000 |
| 2 µg of HaloTag-Myo10-FlagTag, 2 µg of EGFP-DCC ΔP1ΔP3, 1 µg of netrin | Colocalization heatmap | Lipofectamine 3000 |
| 2 µg of HaloTag-Myo10-2mut | Colocalization heatmap | Lipofectamine 3000 |
| 2 µg of HaloTag-Myo10-7mut | Colocalization heatmap | Lipofectamine 3000 |

### Staining, imaging, and analysis for protein colocalization heat maps

After 3-4 h of seeding onto 8-well chambered coverslip, cells were fixed for 20 min in a solution comprising: 4% PFA, 0.08% Triton, 0.5 µM TMR-HaloLigand, and 1:1000 DAPI in PEM buffer (0.1 M PIPES, 5 mM EGTA, 2 mM MgCl_2_·6H_2_O, pH=6.95). Cells were washed 4x with PBS before immediate imaging.

For immunostaining of active beta-1 integrin, cells were fixed as described above. Cells were blocked with 3% BSA in PBS for 25 min at RT in the dark before incubation with anti-integrin antibody (12G10, 500x dilution) in 3% BSA shaking overnight at 4°C in the dark, at 60 rpm. The next morning, cells were washed 3x with PBS (4 min each) before staining with anti-mouse secondary antibody conjugated to AlexaFluor 647 (800x dilution) in 3% BSA in PBS for 50 min at RT in the dark. Cells were washed 3× with PBS (4 min each), post-fixed for 10 min with 4% PFA in PBS, then washed 4x with PBS before immediate imaging.

Fixed cells were imaged at 170 ms exposure and 125 gain using an epifluorescence microscope with 60x, 1.2 NA water objective and LED light source (Zeiss Axiovert 200). Cells expressing both EGFP-DCC and HaloTag-Myo10 positive cells were imaged with blue light (460 nm for EGFP), green light (520 nm for TMR), then red light (630 nm for AF647). Images were saved as a stack in Micro-Manager (Version 2.0).

To map protein localization within filopodia, a modified method developed by Miihkinen et al. was utilized.^14^ Image stacks were first loaded into ImageJ. On the merged image, line intensity profiles (3-pixel width) were manually drawn on the merged image from filopodium tip to base; the membrane protein channel served as a reference for the cell body. Only filopodia with Myo10 signal were analyzed, and crowded, overlapping filopodia were excluded. A zip folder of filopodia traces were saved per cell. Image stacks were background subtracted using a rolling ball radius of 50 pixels and split into separate protein tiff images. An ImageJ macro was then used to loop through all cell images and apply the saved ROI traces to each protein tiff image, saving the results of “distance vs. gray value” as a csv file per cell. The csv files were analyzed in R.

Each filopodium was binned (bins set to 7 or 14 depending on the experiment), with the sum of the bins set to 100 so that all filopodia were normalized for length and signal intensity. The median value in each bin across all filopodia is represented in the heatmaps. Signal at filopodial tips was defined as the first bin of filopodia binned to 7 total bins. Filopodial lengths were extracted from the line intensity profiles. Three bioreplicates per condition were analyzed.

Cargo protein (active integrin, DCC) and Myo10 correlation was defined as the Pearson correlation coefficient of the line intensity profiles of a given filopodium’s Myo10 and cargo signal.

### Live cell TIRF microscopy and live image analysis

After 3-4 h of seeding onto round coverslip, cells were incubated in a TC incubator with 200 µL of 500 µM TMR-HaloLigand in prewarmed DMEM + 10% FBS. After 10 min, cells were gently washed 2× with 200 µL of prewarmed DMEM + 10% FBS and then imaged using a 100x, 1.65 NA objective (Olympus) on a custom-built total internal reflection microscope equipped with an electron-multiplying charge-coupled device (EMCCD) camera (iXon; Andor Technologies). The 100x oil objective was heated to 37°C for 45 min prior to imaging to equilibrate the temperature of the objective. Cells were simultaneously excited with 488 nm and 561 nm lasers, both at 2% power, passing through a dichroic mirror and a Cy3/GFP emission filter for a split screen acquisition of both channels at 200 ms exposure, an electron multiplication gain of 200 and 1s delay between images. Cells were recorded for 5-10 min or until the cell showed signs of retraction/death. Each movie was saved as a stack in Micro-Manager (Version 2.0).

To obtain correlation values for Myo10 and DCC filopodial tip puncta signal in nascent filopodia, image stacks were first loaded into ImageJ. Because the fluorescent probe/protein intensities differ between channels, the left and right halves of the field of view were contrast-adjusted to their own minimum and maximum values, applied to the entire stack, and background subtracted using a 50-pixel rolling ball radius. Newly growing filopodia (i.e., originating from the plasma membrane, not from existing filopodia) were then manually tracked using the “Manual Tracking” plugin with “local barycentre” centering correction and an 8-pixel search square. The start of trajectories were marked 2-10 frames before a punctum at the plasma membrane moved away from the cell body into a new filopodium. Signal-over-time data were saved as csv files per cell and imported into R to calculate the Pearson correlation coefficient between Myo10 and cargo signal traces for each filopodium. Four bioreplicates per condition were analyzed.

### Staining of DNA-PAINT samples

After 5 h of seeding onto 8-well chambered coverslip, cells were fixed in pre-warmed 4% PFA in PBS for 15 min. Cells were washed 3x with PBS and incubated for 5 min in a 2x dilution of 90 nm gold nanoparticles in PBS. After 3x PBS washes, the cells were blocked/permeabilized in PBS containing 3% BSA and 0.2% Triton X-100 for 30 min. After 3x PBS washes, the cells were stained with the following antibodies at 4 °C overnight in 200 µL of antibody incubation buffer (PBS containing 1 mM EDTA, 0.02% Tween-20, 0.05% NaN3, 2% BSA, 0.05 mg/mL sheared salmon sperm DNA): Halo-5xR2 (1 µM); 25 nM of GFP nanobody clones 1H1 and 1B2 conjugated to 5xR1 (nanobodies from Nanotag, cat. no. N0305); mouse anti-human CD29 clone 4B7R (cat. no. MAB6944, 1 µg/µL) at 1:200 dilution; 25 nM of sdAb anti-mouse IgG1 clone 10A4 (nanotag, cat. no. N2005) conjugated to DNA docking strand 7xR3. Full DNA sequences for these constructs were previously reported.^58^ The next morning, cells were quickly washed 3x with PBS, left in Buffer C (PBS, 1 mM EDTA, 500 mM NaCl pH 7.4, 0.02% Tween) for 10 min, quickly washed 3x with PBS, post-fixed for 10 min with 4% PFA in PBS, and washed 3x with PBS.

Samples were imaged in buffer C supplemented with 1× trolox and target protein imager strand (around 100 pM, depending on localization density) conjugated to Cy3B: R1 for DCC, R2 for Myo10, and R3 for integrin. Cells were typically imaged for 30,000-40,000 frames with 100 ms exposure at 40-50 mW laser power measured after the objective, where 45 mW corresponds to a power density of 225 W cm^−2^. Targets were imaged sequentially, by which the sample was washed with buffer C between targets until undetectable signal from the previous imager solution. All DNA-PAINT labeling reagents were kindly supplied by Jungmann Lab.

Seven cells were imaged for integrin and Myo10 samples (24 filopodial regions analyzed). Ten cells were imaged for DCC and Myo10 samples (35 filopodial regions analyzed).

### DNA-PAINT imaging and microscopy set-up

Imaging was conducted on an inverted microscope (Nikon Instruments, Eclipse Ti2) with the Perfect Focus System and an oil-immersion objective (Nikon Instruments, Apo SR TIRF ×100, NA 1.49). 488 nm and 560 (MPB Communications, 1 W each) were used for excitation and coupled into the microscope via a Nikon manual TIRF module. Laser beams were passed through cleanup filters (Chroma Technology, ZET488/10x for 488 nm, ZET561/10x for 560 nm) and coupled into the objective with a beam splitter (Chroma Technology, ZT488rdc-UF2 for 488 nm, ZT561rdc-UF2 for 560 nm). Fluorescence was spectrally filtered with emission filters (Chroma Technology, ET525/50m and ET500lp for 488 nm, ET600/50m and ET575lp for 560 nm) and imaged on an sCMOS camera (Hamamatsu, ORCA-Fusion BT), yielding an effective pixel size of 130 nm after 2×2 binning. 3D imaging was achieved with an astigmatism lens (Nikon Instruments, N-STORM) in the detection path. Raw data was acquired using Micro-Manager (Version 2.0.1).

### DNA-PAINT image analysis

Raw fluorescence single-molecule data were processed using the Picasso software. Single molecules were localized in Picasso using the Gaussian least squares option and a 3D calibration file based on astigmatism ^51^. Reconstructed images were undrifted with gold nanoparticle fiducials. Two-color channels were aligned with cross-correlation of gold nanoparticle fiducials present in both channels. Regions along the filopodia with medium Myo10 and cargo signal intensity were manually picked as circular regions with a 0.6-2.2 µM diameter for downstream analysis and plotting in a custom Jupyter Notebook.

The lowest and highest 5% localization values in x, y and z dimensions were filtered out. Filtered localizations in the picked filopodial regions of interest were clustered by HDBSCAN (minimum samples and cluster size set to 6) to find protein cluster centers in a custom Jupyter Notebook. Cluster centers were used as proxies for single protein positions.

We used PCA to reorient filopodial Myo10 and cargo molecule positions on a common coordinate system. First, we subtracted z-offsets using the mean of the Myo10 localizations. We then performed PCA on the x and y coordinates, rotating the data so that the filopodial axis is aligned to PC1. The tip-base orientation is arbitrary using this procedure, so we tested if the smallest PC1 point was farther than the largest PC1 point from the cell center (the mean of all localizations for that cell). If not, we rotated the xy data about the z-axis by π. To convert protein positions into a polar coordinate system, positions were projected onto the plane orthogonal to the filopodium’s principal axis (PC1), with radius 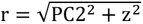 (in nm) and theta 𝜃 = atan2(z, PC2) (in radians).

The local concentrations of Myo10 vs. cargo in the filopodial shaft were estimated by the number of protein molecules within the volume of a cylinder (µm^3^). We started from the PCA-based common coordinate system as described above. For volume calculations, the length of the longitudinal axis (i.e., longest side of the filopodial region, PC1) was used as the height, and the average of half the lengths of the sagittal (PC2) and the transverse axis (PC3) was used as the radius. Protein molecules per µm^3^ was converted to µM.

To calculate percent of ligand bound (%L_bound_ = PL/LT), the following equation was used^59^: 𝑃𝐿 = ([𝐾𝐷 + 𝐿𝑇 + 𝑃𝑇] − √((𝐾𝐷 + 𝐿𝑇 + 𝑃𝑇)² − 4(𝑃𝑇)(𝐿𝑇))) / 2, where PT = total Myo10 concentration, LT = total cargo protein concentration, PL = Myo10-cargo complex concentration, and the K_D_ values reported from literature.

### HaloTag labeling efficiency by DNA-PAINT

After seeding 20,000 CHO-K1 WT cells per well in Ibidi µ-Slide 8 Well^high^, cells were transiently transfected (after ≥4 h) with mEGFP-CD86-HaloTag plasmid using Lipofectamine 3000. The following day, cells were fixed with pre-warmed 4% PFA in PBS for 15 min, washed 3x with PBS, and permeabilized/blocked for 30 min in antibody incubation buffer containing 0.2% Triton X-100. Then, cells were incubated overnight with Halo-5xR2 (1 µM) and GFP nanobody clone 1H1 -7xR4 (25 nM) in antibody incubation buffer; untransfected control cells received only Halo-5xR2. Samples were washed 3x with PBS, incubated in Buffer C for 10 min, post-fixed (4% PFA, 10 min), washed 3x, incubated with undiluted gold nanoparticles for 5 min, and washed 3x. Imaging was performed in Buffer C with 400 pM imager strands (R2, R4), acquiring ∼40,000 frames at 100 ms exposure and 30 mW laser power.

Single-molecule data were undrifted, aligned, and filtered by sx, sy, and net gradient. Proteins were identified from localization clusters in Picasso using the function “SMLM clusterer” (radius 8 nm and min 10 localizations, with “basic frame analysis” to exclude imager sticking events), and homogeneous regions were selected for analysis. Labeling efficiency was quantified in Picasso Spinna (v0.9.9) using a 12 nm GFP–Halo distance, 5 nm localization uncertainty and a granularity of 101 for the parameter search space.

## References

(1) Berg, J. S.; Derfler, B. H.; Pennisi, C. M.; Corey, D. P.; Cheney, R. E. Myosin-X, a Novel Myosin with Pleckstrin Homology Domains, Associates with Regions of Dynamic Actin. Journal of Cell Science 2000, 113 (19), 3439–3451. 10.1242/jcs.113.19.3439.

(2) Kerber, M. L.; Cheney, R. E. Myosin-X: A MyTH-FERM Myosin at the Tips of Filopodia. Journal of Cell Science 2011, 124 (22), 3733–3741. 10.1242/jcs.023549.

(3) Nagy, S.; Ricca, B. L.; Norstrom, M. F.; Courson, D. S.; Brawley, C. M.; Smithback, P. A.; Rock, R. S. A Myosin Motor That Selects Bundled Actin for Motility. Proc. Natl. Acad. Sci. U.S.A. 2008, 105 (28), 9616–9620. 10.1073/pnas.0802592105.

(4) Ropars, V.; Yang, Z.; Isabet, T.; Blanc, F.; Zhou, K.; Lin, T.; Liu, X.; Hissier, P.; Samazan, F.; Amigues, B.; Yang, E. D.; Park, H.; Pylypenko, O.; Cecchini, M.; Sindelar, C. V.; Sweeney, H. L.; Houdusse, A. The Myosin X Motor Is Optimized for Movement on Actin Bundles. Nat Commun 2016, 7 (1), 12456. 10.1038/ncomms12456.

(5) Petersen, K. J.; Goodson, H. V.; Arthur, A. L.; Luxton, G. W. G.; Houdusse, A.; Titus, M. A. MyTH4-FERM Myosins Have an Ancient and Conserved Role in Filopod Formation. Proc. Natl. Acad. Sci. U.S.A. 2016, 113 (50). 10.1073/pnas.1615392113.

(6) Arthur, A. L.; Songster, L. D.; Sirkia, H.; Bhattacharya, A.; Kikuti, C.; Borrega, F. P.; Houdusse, A.; Titus, M. A. Optimized Filopodia Formation Requires Myosin Tail Domain Cooperation. Proc. Natl. Acad. Sci. U.S.A. 2019, 116 (44), 22196–22204. 10.1073/pnas.1901527116.

(7) Houdusse, A.; Titus, M. A. The Many Roles of Myosins in Filopodia, Microvilli and Stereocilia. Current Biology 2021, 31 (10), R586–R602. 10.1016/j.cub.2021.04.005.

(8) Mattila, P. K.; Lappalainen, P. Filopodia: Molecular Architecture and Cellular Functions. Nat Rev Mol Cell Biol 2008, 9 (6), 446–454. 10.1038/nrm2406.

(9) Tokuo, H.; Ikebe, M. Myosin X Transports Mena/VASP to the Tip of Filopodia. Biochemical and Biophysical Research Communications 2004, 319 (1), 214–220. 10.1016/j.bbrc.2004.04.167.

(10) Weber, K. L.; Sokac, A. M.; Berg, J. S.; Cheney, R. E.; Bement, W. M. A Microtubule-Binding Myosin Required for Nuclear Anchoring and Spindle Assembly. Nature 2004, 431 (7006), 325–329. 10.1038/nature02834.

(11) Hirano, Y.; Hatano, T.; Takahashi, A.; Toriyama, M.; Inagaki, N.; Hakoshima, T. Structural Basis of Cargo Recognition by the Myosin-X MyTH4-FERM Domain: Myosin-X Binding to DCC, Integrin and Microtubule. The EMBO Journal 2011, 30 (13), 2734–2747. 10.1038/emboj.2011.177.

(12) Pi, X.; Ren, R.; Kelley, R.; Zhang, C.; Moser, M.; Bohil, A. B.; DiVito, M.; Cheney, R. E.; Patterson, C. Sequential Roles for Myosin-X in BMP6-Dependent Filopodial Extension, Migration, and Activation of BMP Receptors. The Journal of Cell Biology 2007, 179 (7), 1569–1582. 10.1083/jcb.200704010.

(13) Almagro, S.; Durmort, C.; Chervin-Pétinot, A.; Heyraud, S.; Dubois, M.; Lambert, O.; Maillefaud, C.; Hewat, E.; Schaal, J. P.; Huber, P.; Gulino-Debrac, D. The Motor Protein Myosin-X Transports VE-Cadherin along Filopodia To Allow the Formation of Early Endothelial Cell-Cell Contacts. Molecular and Cellular Biology 2010, 30 (7), 1703–1717. 10.1128/MCB.01226-09.

(14) Miihkinen, M.; Grönloh, M. L. B.; Popović, A.; Vihinen, H.; Jokitalo, E.; Goult, B. T.; Ivaska, J.; Jacquemet, G. Myosin-X and Talin Modulate Integrin Activity at Filopodia Tips. Cell Reports 2021, 36 (11), 109716. 10.1016/j.celrep.2021.109716.

(15) Wei, Z.; Yan, J.; Lu, Q.; Pan, L.; Zhang, M. Cargo Recognition Mechanism of Myosin X Revealed by the Structure of Its Tail MyTH4-FERM Tandem in Complex with the DCC P3 Domain. Proc Natl Acad Sci USA 2011, 108 (9), 3572–3577. 10.1073/pnas.1016567108.

(16) Liu, Y.; Peng, Y.; Dai, P.-G.; Du, Q.-S.; Mei, L.; Xiong, W.-C. Differential Regulation of Myosin X Movements by Its Cargos, DCC and Neogenin. Journal of Cell Science 2012, 125 (3), 751–762. 10.1242/jcs.094946.

(17) Hynes, R. O. Integrins. Cell 2002, 110 (6), 673–687. 10.1016/S0092-8674(02)00971-6.

(18) Pfaff, M.; Liu, S.; Erle, D. J.; Ginsberg, M. H. Integrin β Cytoplasmic Domains Differentially Bind to Cytoskeletal Proteins. Journal of Biological Chemistry 1998, 273 (11), 6104–6109. 10.1074/jbc.273.11.6104.

(19) Luo, B.-H.; Carman, C. V.; Springer, T. A. Structural Basis of Integrin Regulation and Signaling. Annu. Rev. Immunol. 2007, 25 (1), 619–647. 10.1146/annurev.immunol.25.022106.141618.

(20) Zhang, H.; Berg, J. S.; Li, Z.; Wang, Y.; Lång, P.; Sousa, A. D.; Bhaskar, A.; Cheney, R. E.; Strömblad, S. Myosin-X Provides a Motor-Based Link between Integrins and the Cytoskeleton. Nat Cell Biol 2004, 6 (6), 523–531. 10.1038/ncb1136.

(21) Arjonen, A.; Kaukonen, R.; Mattila, E.; Rouhi, P.; Högnäs, G.; Sihto, H.; Miller, B. W.; Morton, J. P.; Bucher, E.; Taimen, P.; Virtakoivu, R.; Cao, Y.; Sansom, O. J.; Joensuu, H.; Ivaska, J. Mutant P53–Associated Myosin-X Upregulation Promotes Breast Cancer Invasion and Metastasis. J. Clin. Invest. 2014, 124 (3), 1069–1082. 10.1172/JCI67280.

(22) Courson, D. S.; Cheney, R. E. Myosin-X and Disease. Experimental Cell Research 2015, 334 (1), 10–15. 10.1016/j.yexcr.2015.03.014.

(23) Zhu, X.-J.; Wang, C.-Z.; Dai, P.-G.; Xie, Y.; Song, N.-N.; Liu, Y.; Du, Q.-S.; Mei, L.; Ding, Y.-Q.; Xiong, W.-C. Myosin X Regulates Netrin Receptors and Functions in Axonal Path-Finding. Nat Cell Biol 2007, 9 (2), 184–192. 10.1038/ncb1535.

(24) Yu, H.-L.; Peng, Y.; Zhao, Y.; Lan, Y.-S.; Wang, B.; Zhao, L.; Sun, D.; Pan, J.-X.; Dong, Z.-Q.; Mei, L.; Ding, Y.-Q.; Zhu, X.-J.; Xiong, W.-C. Myosin X Interaction with KIF13B, a Crucial Pathway for Netrin-1-Induced Axonal Development. J. Neurosci. 2020, 40 (48), 9169–9185. 10.1523/JNEUROSCI.0929-20.2020.

(25) Shibata, D.; Reale, M. A.; Lavin, P.; Silverman, M.; Fearon, E. R.; Steele, G.; Jessup, J. M.; Loda, M.; Summerhayes, I. C. The DCC Protein and Prognosis in Colorectal Cancer. N Engl J Med 1996, 335 (23), 1727–1732. 10.1056/NEJM199612053352303.

(26) Anthis, N. J.; Wegener, K. L.; Ye, F.; Kim, C.; Goult, B. T.; Lowe, E. D.; Vakonakis, I.; Bate, N.; Critchley, D. R.; Ginsberg, M. H.; Campbell, I. D. The Structure of an Integrin/Talin Complex Reveals the Basis of inside-out Signal Transduction. EMBO J 2009, 28 (22), 3623–3632. 10.1038/emboj.2009.287.

(27) García-Alvarez, B.; de Pereda, J. M.; Calderwood, D. A.; Ulmer, T. S.; Critchley, D.; Campbell, I. D.; Ginsberg, M. H.; Liddington, R. C. Structural Determinants of Integrin Recognition by Talin. Molecular Cell 2003, 11 (1), 49–58. 10.1016/S1097-2765(02)00823-7.

(28) Kolodziej, P. A.; Timpe, L. C.; Mitchell, K. J.; Fried, S. R.; Goodman, C. S.; Jan, L. Y.; Jan, Y. N. Frazzled Encodes a Drosophila Member of the DCC Immunoglobulin Subfamily and Is Required for CNS and Motor Axon Guidance. Cell 1996, 87 (2), 197– 204. 10.1016/S0092-8674(00)81338-0.

(29) Holehouse, A. S.; Kragelund, B. B. The Molecular Basis for Cellular Function of Intrinsically Disordered Protein Regions. Nat Rev Mol Cell Biol 2024, 25 (3), 187–211. 10.1038/s41580-023-00673-0.

(30) Tompa, P.; Fuxreiter, M. Fuzzy Complexes: Polymorphism and Structural Disorder in Protein–Protein Interactions. Trends in Biochemical Sciences 2008, 33 (1), 2–8. 10.1016/j.tibs.2007.10.003.

(31) Zhou, H.-X. Intrinsic Disorder: Signaling via Highly Specific but Short-Lived Association. Trends in Biochemical Sciences 2012, 37 (2), 43–48. 10.1016/j.tibs.2011.11.002.

(32) Fuxreiter, M. Classifying the Binding Modes of Disordered Proteins. IJMS 2020, 21 (22), 8615. 10.3390/ijms21228615.

(33) Englander, S. W. Hydrogen Exchange and Mass Spectrometry: A Historical Perspective. J. Am. Soc. Mass Spectrom. 2006, 17 (11), 1481–1489. 10.1016/j.jasms.2006.06.006.

(34) Mayne, L. Hydrogen Exchange Mass Spectrometry. In Methods in Enzymology; Elsevier, 2016; Vol. 566, pp 335–356. 10.1016/bs.mie.2015.06.035.

(35) Hamuro, Y. Tutorial: Chemistry of Hydrogen/Deuterium Exchange Mass Spectrometry. J. Am. Soc. Mass Spectrom. 2021, 32 (1), 133–151. 10.1021/jasms.0c00260.

(36) Lin, X.; Molina, A. V.; Shangguan, J.; Chen, R.; Sosnick, T. R. Practical Tips for the Application of HDX-MS to Membrane Proteins, Biomolecular Condensates, and Weak Protein Binders. J. Am. Soc. Mass Spectrom. 2025, 36 (8), 1575–1587. 10.1021/jasms.5c00067.

(37) Castets, M.; Broutier, L.; Molin, Y.; Brevet, M.; Chazot, G.; Gadot, N.; Paquet, A.; Mazelin, L.; Jarrosson-Wuilleme, L.; Scoazec, J.-Y.; Bernet, A.; Mehlen, P. DCC Constrains Tumour Progression via Its Dependence Receptor Activity. Nature 2012, 482 (7386), 534–537. 10.1038/nature10708.

(38) Mehlen, P.; Rabizadeh, S.; Snipas, S. J.; Assa-Munt, N.; Salvesen, G. S.; Bredesen, D. E. The DCC Gene Product Induces Apoptosis by a Mechanism Requiring Receptor Proteolysis. Nature 1998, 395 (6704), 801–804. 10.1038/27441.

(39) Hirano, Y.; Hatano, T.; Takahashi, A.; Toriyama, M.; Inagaki, N.; Hakoshima, T. Structural Basis of Cargo Recognition by the Myosin-X MyTH4-FERM Domain: Myosin-X Binding to DCC, Integrin and Microtubule. The EMBO Journal 2011, 30 (13), 2734–2747. 10.1038/emboj.2011.177.

(40) Endesfelder, U.; Malkusch, S.; Fricke, F.; Heilemann, M. A Simple Method to Estimate the Average Localization Precision of a Single-Molecule Localization Microscopy Experiment. Histochem Cell Biol 2014, 141 (6), 629–638. 10.1007/s00418-014-1192-3.

(41) Campello, R. J. G. B.; Moulavi, D.; Sander, J. Density-Based Clustering Based on Hierarchical Density Estimates. In Advances in Knowledge Discovery and Data Mining; Pei, J., Tseng, V. S., Cao, L., Motoda, H., Xu, G., Eds.; Hutchison, D., Kanade, T., Kittler, J., Kleinberg, J. M., Mattern, F., Mitchell, J. C., Naor, M., Nierstrasz, O., Pandu Rangan, C., Steffen, B., Sudan, M., Terzopoulos, D., Tygar, D., Vardi, M. Y., Weikum, G., Series Eds.; Lecture Notes in Computer Science; Springer Berlin Heidelberg: Berlin, Heidelberg, 2013; Vol. 7819, pp 160–172. 10.1007/978-3-642-37456-2_14.

(42) Hellmeier, J.; Strauss, S.; Xu, S.; Masullo, L. A.; Unterauer, E. M.; Kowalewski, R.; Jungmann, R. Quantification of Absolute Labeling Efficiency at the Single-Protein Level. Nat Methods 2024, 21 (9), 1702–1707. 10.1038/s41592-024-02242-5.

(43) Nagy, S.; Rock, R. S. Structured Post-IQ Domain Governs Selectivity of Myosin X for Fascin-Actin Bundles. Journal of Biological Chemistry 2010, 285 (34), 26608–26617. 10.1074/jbc.M110.104661.

(44) Liu, K. C.; Jacobs, D. T.; Dunn, B. D.; Fanning, A. S.; Cheney, R. E. Myosin-X Functions in Polarized Epithelial Cells. MBoC 2012, 23 (9), 1675–1687. 10.1091/mbc.e11-04-0358.

(45) Iešmantavičius, V.; Dogan, J.; Jemth, P.; Teilum, K.; Kjaergaard, M. Helical Propensity in an Intrinsically Disordered Protein Accelerates Ligand Binding. Angew. Chem. Int. Ed. 2014, 53 (6), 1548–1551. 10.1002/anie.201307712.

(46) Moses, D.; Guadalupe, K.; Yu, F.; Flores, E.; Perez, A.; McAnelly, R.; Shamoon, N. M.; Cuevas-Zepeda, E.; Merg, A. D.; Martin, E. W.; Holehouse, A. S.; Sukenik, S. Structural Biases in Disordered Proteins Are Prevalent in the Cell; preprint; Biophysics, 2021. 10.1101/2021.11.24.469609.

(47) Elkjær, S.; Due, A. D.; Christensen, L. F.; Theisen, F. F.; Staby, L.; Kragelund, B. B.; Skriver, K. Evolutionary Fine-Tuning of Residual Helix Structure in Disordered Proteins Manifests in Complex Structure and Lifetime. Commun Biol 2023, 6 (1), 63. 10.1038/s42003-023-04445-6.

(48) Roca-Cusachs, P.; Gauthier, N. C.; Del Rio, A.; Sheetz, M. P. Clustering of α_5_ β_1_ Integrins Determines Adhesion Strength Whereas α_v_ β_3_ and Talin Enable Mechanotransduction. Proc. Natl. Acad. Sci. U.S.A. 2009, 106 (38), 16245–16250. 10.1073/pnas.0902818106.

(49) Friedland, J. C.; Lee, M. H.; Boettiger, D. Mechanically Activated Integrin Switch Controls α_5_ β_1_ Function. Science 2009, 323 (5914), 642–644. 10.1126/science.1168441.

(50) Arias-Salgado, E. G.; Lizano, S.; Sarkar, S.; Brugge, J. S.; Ginsberg, M. H.; Shattil, S. J. Src Kinase Activation by Direct Interaction with the Integrin β Cytoplasmic Domain. Proc. Natl. Acad. Sci. U.S.A. 2003, 100 (23), 13298–13302. 10.1073/pnas.2336149100.

(51) Anthis, N. J.; Haling, J. R.; Oxley, C. L.; Memo, M.; Wegener, K. L.; Lim, C. J.; Ginsberg, M. H.; Campbell, I. D. β Integrin Tyrosine Phosphorylation Is a Conserved Mechanism for Regulating Talin-Induced Integrin Activation. Journal of Biological Chemistry 2009, 284 (52), 36700–36710. 10.1074/jbc.M109.061275.

(52) Shangguan, J.; Rock, R. S. Hundreds of Myosin 10s Are Pushed to the Tips of Filopodia and Could Cause Traffic Jams on Actin. eLife 2024, 12, RP90603. 10.7554/eLife.90603.4.

(53) Courson, D. S.; Rock, R. S. Actin Cross-Link Assembly and Disassembly Mechanics for α-Actinin and Fascin. Journal of Biological Chemistry 2010, 285 (34), 26350–26357. 10.1074/jbc.M110.123117.

(54) Xu, S.; Liu, Y.; Li, X.; Liu, Y.; Meijers, R.; Zhang, Y.; Wang, J. The Binding of DCC-P3 Motif and FAK-FAT Domain Mediates the Initial Step of Netrin-1/DCC Signaling for Axon Attraction. Cell Discov 2018, 4 (1), 8. 10.1038/s41421-017-0008-8.

(55) Ren, X.; Ming, G.; Xie, Y.; Hong, Y.; Sun, D.; Zhao, Z.; Feng, Z.; Wang, Q.; Shim, S.; Chen, Z.; Song, H.; Mei, L.; Xiong, W. Focal Adhesion Kinase in Netrin-1 Signaling. Nat Neurosci 2004, 7 (11), 1204–1212. 10.1038/nn1330.

(56) Chen, H.-C.; Appeddu, P. A.; Parsons, J. T.; Hildebrand, J. D.; Schaller, M. D.; Guan, J.-L. Interaction of Focal Adhesion Kinase with Cytoskeletal Protein Talin. Journal of Biological Chemistry 1995, 270 (28), 16995–16999. 10.1074/jbc.270.28.16995.

(57) Kanchanawong, P.; Shtengel, G.; Pasapera, A. M.; Ramko, E. B.; Davidson, M. W.; Hess, H. F.; Waterman, C. M. Nanoscale Architecture of Integrin-Based Cell Adhesions. Nature 2010, 468 (7323), 580–584. 10.1038/nature09621.

(58) Strauss, S.; Jungmann, R. Up to 100-Fold Speed-up and Multiplexing in Optimized DNA-PAINT. Nat Methods 2020, 17 (8), 789–791. 10.1038/s41592-020-0869-x.

(59) Kochert, B. A.; Iacob, R. E.; Wales, T. E.; Makriyannis, A.; Engen, J. R. Hydrogen-Deuterium Exchange Mass Spectrometry to Study Protein Complexes. In Protein Complex Assembly: Methods and Protocols; Marsh, J. A., Ed.; Methods in Molecular Biology; Springer: New York, NY, 2018; pp 153–171. 10.1007/978-1-4939-7759-8_10.

