## Supplemental Figures for "Dynamic Myosin 10 coupling to DCC and β1 integrin is mediated by intrinsically disordered regions during filopodial transport and patterning"

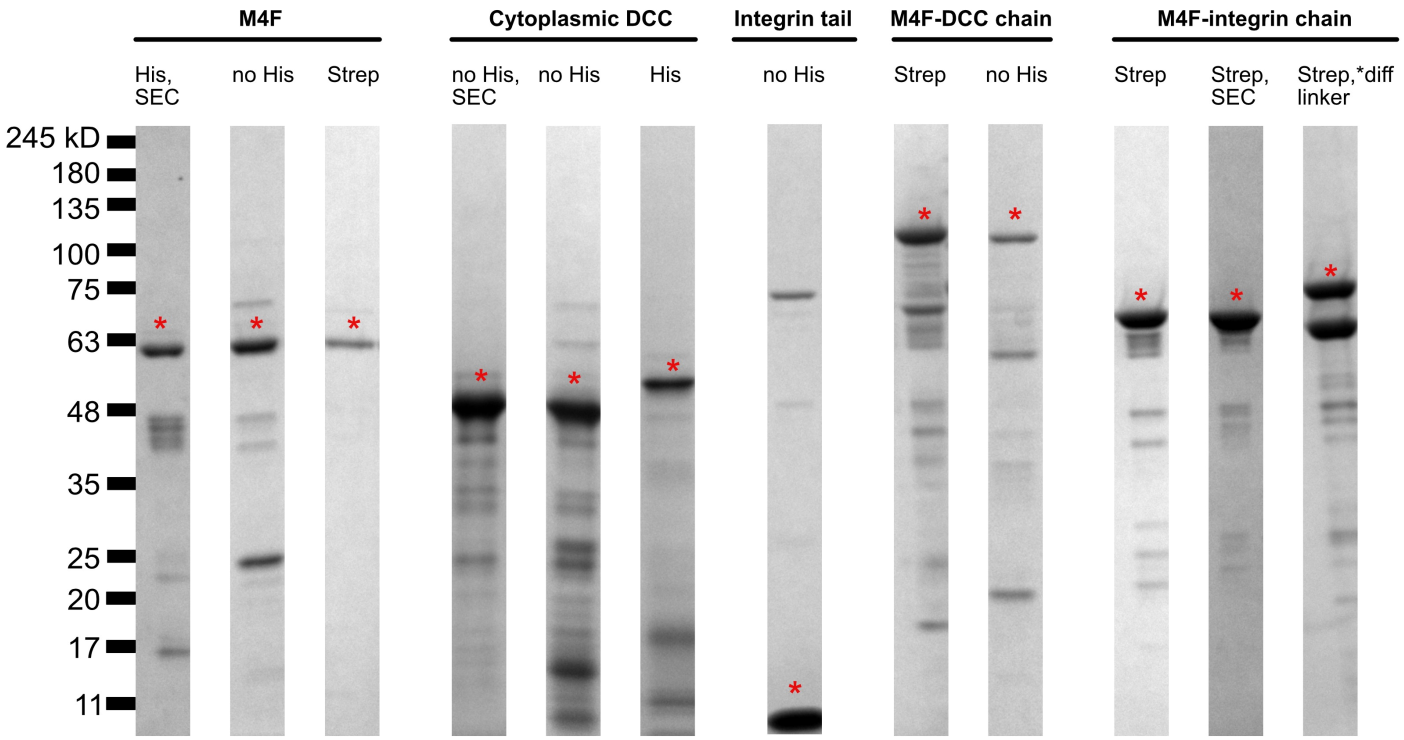


**Supplementary Figure 1. SDS-PAGE gels of the protein bioreplicates used for the HDX-MS experiments.** A red asterisk is marked above the protein of interest.

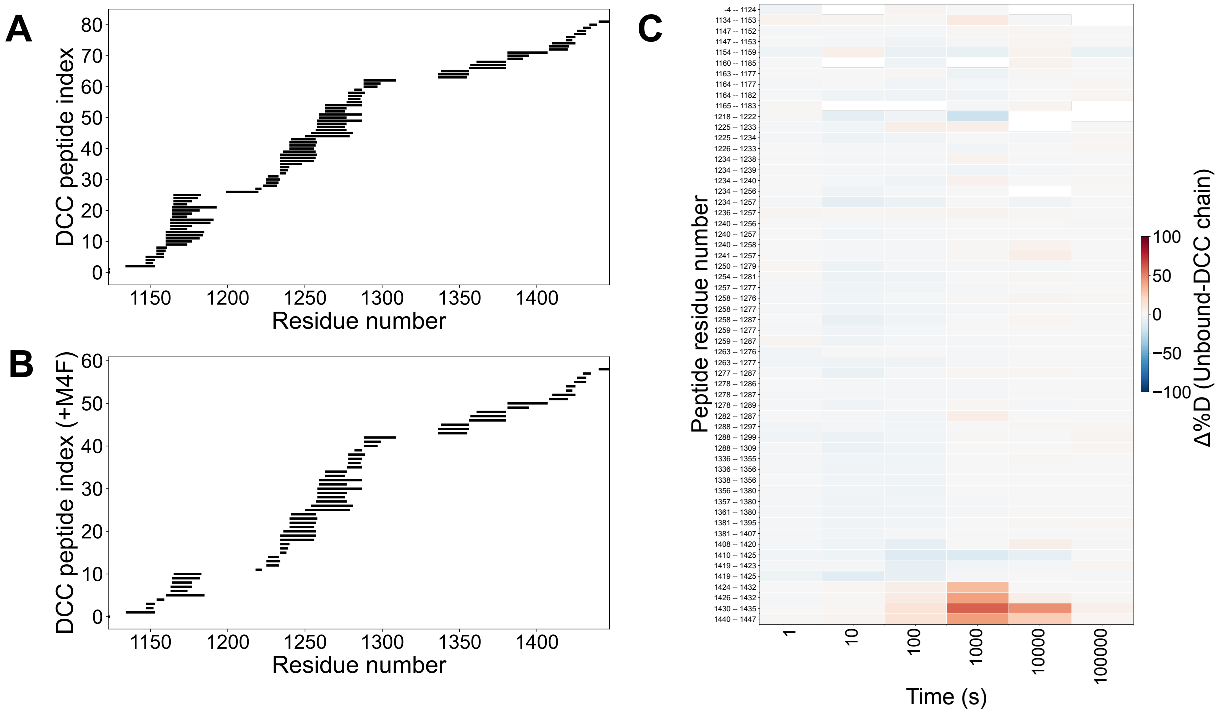


**Supplementary Figure 2. HDX-MS sequence coverage of cytoplasmic DCC and deuteration differences of upon Myo10 M4F binding.** Peptide sequence coverage for A) unbound cytoplasmic DCC, B) cytoplasmic DCC bound to M4F (tethered construct). The residue numbers correspond to the position in the full-length protein sequences. C) Heatmap showing deuteration changes for peptides detected in both unbound and M4F-bound cytoplasmic DCC conditions across labeling time points. Red indicates less deuteration upon binding. Peptide residue ranges are displayed on the y-axis.

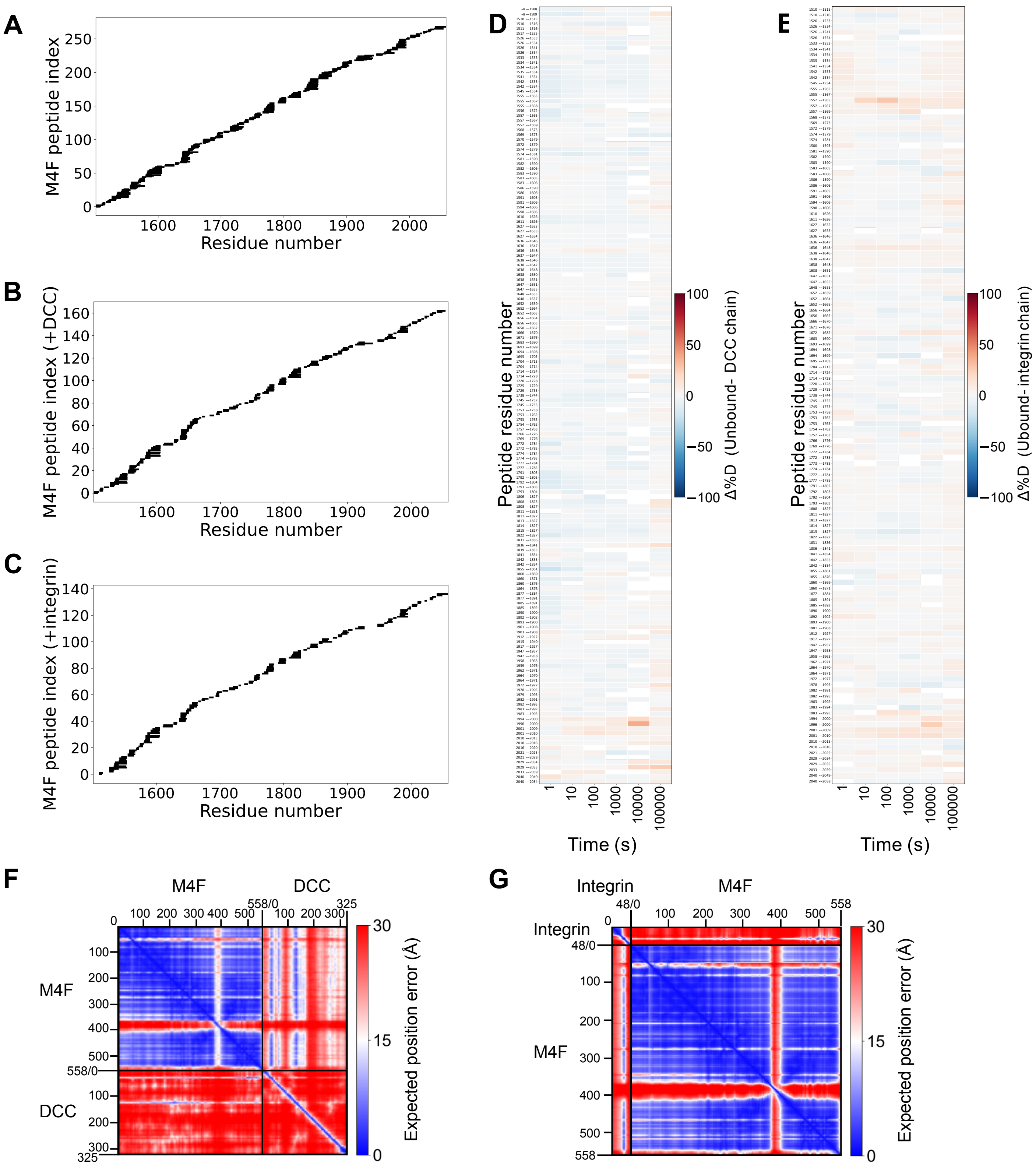


**Supplementary Figure 3. HDX-MS sequence coverage of Myo10 M4F and deuteration differences of upon cargo binding.** Peptide sequence coverage for A) unbound M4F, B) M4F bound to cytoplasmic DCC (tethered construct), C) M4F bound to cytoplasmic β1 integrin (tethered construct). The residue numbers correspond to the position in the full-length protein sequences. D) Heatmap showing deuteration changes for peptides detected in both unbound and cytoplasmic DCC-bound M4F conditions across labeling time points. E) Heatmap showing deuteration changes for peptides detected in both unbound and cytoplasmic β1 integrin -bound M4F conditions across labeling time points. For D & E, red indicates less deuteration upon binding. Peptide residue ranges are displayed on the y-axis. F) PAE plot of the AlphaFold-Multimer structure of M4F and cytoplasmic DCC. G) PAE plot of the AlphaFold3 structure of M4F and cytoplasmic β1 integrin.

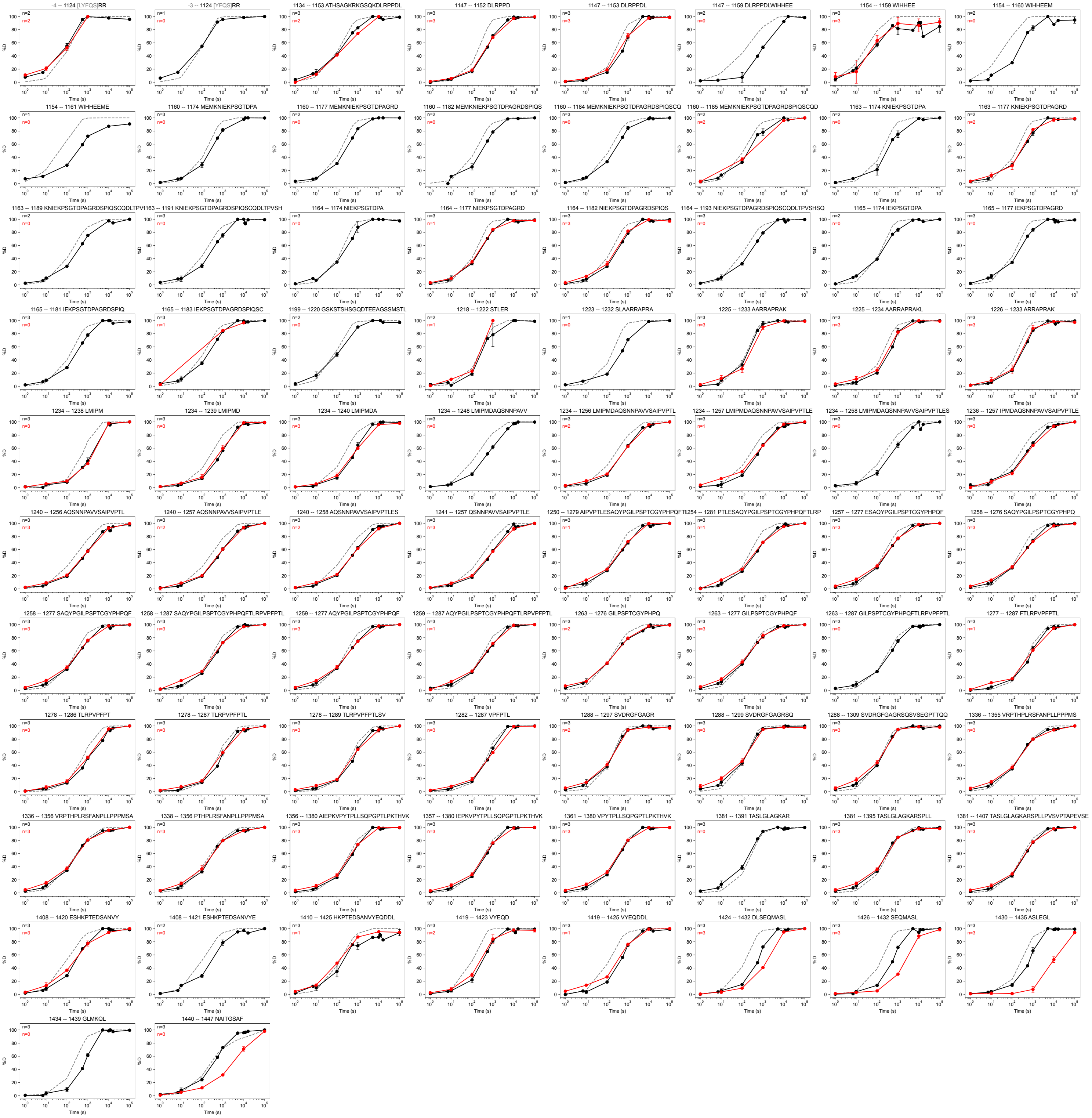


**Supplementary Figure 4. Deuterium uptake plots for all cytoplasmic DCC peptides.** Unbound DCC = black line, bound to M4F (single-chain construct) = red line, $k_{chem}$ = gray dashed line. Numbers above each plot correspond to the residue positions on the full-length DCC sequence. Grayed numbers do not belong to the protein sequence (i.e., purification tag). Top left corner states the number of bioreplicates per condition. Data were normalized using in-exchange (0% D) and fully deuterated (100% D) controls; control data points are not shown. Points are means ±SD.

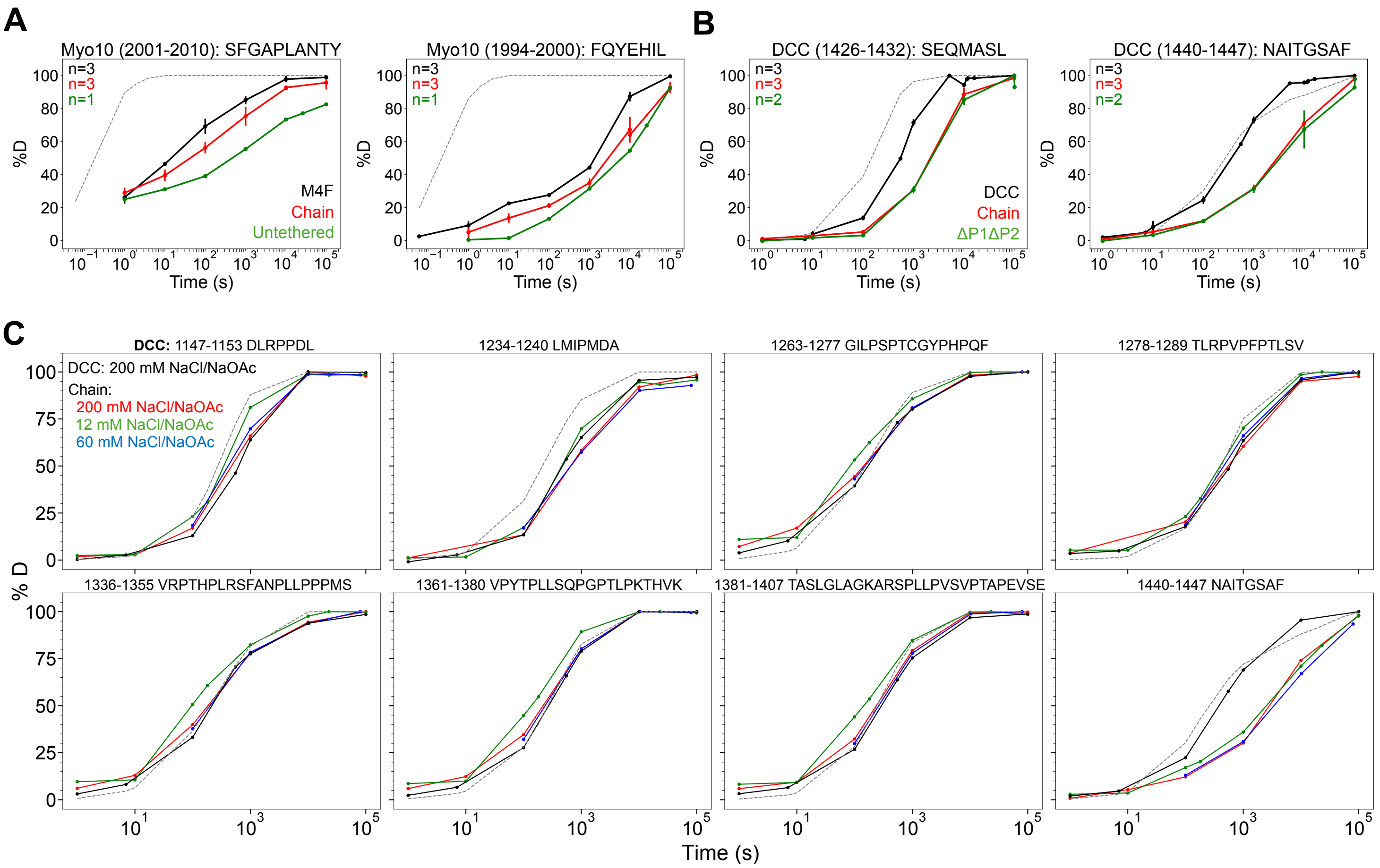


**Supplementary Figure 5. Untethered protein complex has similar HDX pattern as the single-chain construct. Electrostatic interactions are unlikely to be driving binding between cytoplasmic DCC and Myo10 M4F.** A) Non-tethered M4F and cytoplasmic DCC in solution show protection in same M4F sites during binding. Complex was at 20 µM during HDX labeling. No coverage obtained for peptide residues 2029-2035. There was no detectable protection for any cytoplasmic DCC peptides, so that is why the single-chain construct was used for HDX-MS experiments. Unbound M4F = black line, single-chain construct = red line, untethered complex = green line. Numbers above each plot correspond to the residue positions on the full-length Myo10 sequence. Top left corner states the number of bioreplicates per condition. Data were normalized using in-exchange (0% D) and fully deuterated (100% D) controls; control data points are not shown. Points are means ±SD. B) M4F-DCCΔP1ΔP2 single-chain construct shows same protection in P3 motif during binding. Unbound DCC = black line, chain construct = red line, M4F-DCCΔP1ΔP2 = green line. Numbers above each plot correspond to the residue positions on the full-length DCC sequence. Top left corner states the number of bioreplicates per condition. Data were normalized using in-exchange (0% D) and fully deuterated (100% D) controls; control data points are not shown. Points are means ±SD. C) The deuterium uptake plots represent example peptides across cytoplasmic DCC. Numbers above each plot correspond to the residue positions on the full-length DCC sequence. Decreasing salt concentration in the deuterium labeling buffer does not change the observed protection in the presence of M4F. "Bound" state uses single-chain construct. Unbound DCC = black line, bound in 200 mM NaCl/NaOAc (150 mM NaCl, 50 mM NaOAc) = red line, bound in 60 mM NaCl/NaOAc (10 mM NaCl, 50 mM NaOAc) = blue line, bound in 12 mM NaCl/NaOAc (2 mM NaCl, 10 mM NaOAc) = green line, $k_{chem}$ = gray dashed line. 1 bioreplicate displayed per condition.

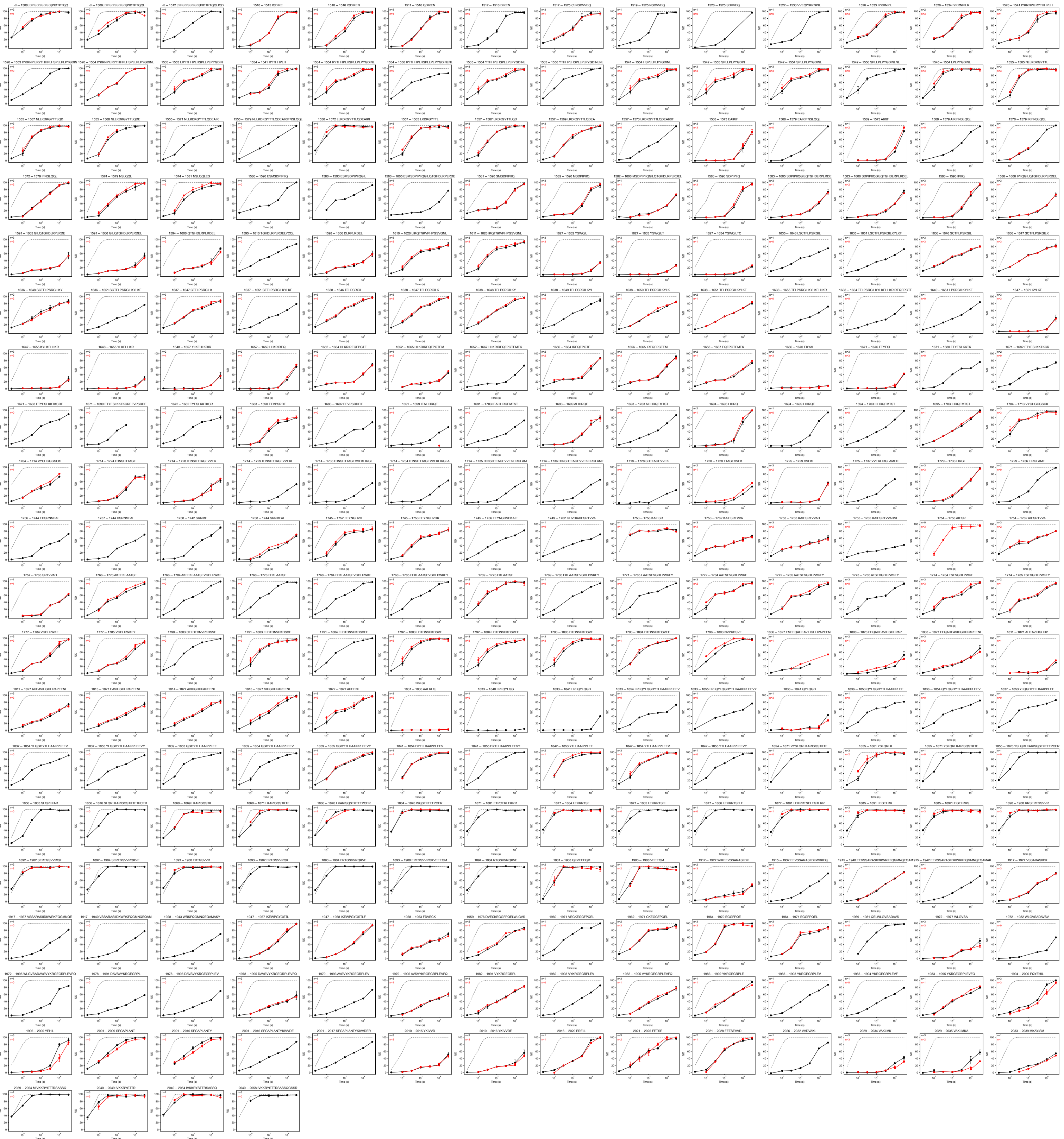

**Supplementary Figure 6. Deuterium uptake plots for all Myo10 M4F peptides.** Unbound M4F = black line, bound to DCC (single-chain construct) = red line, $k_{chem}$ = gray dashed line. Numbers above each plot correspond to the residue positions on the full-length Myo10 sequence. Grayed numbers do not belong to the protein sequence (i.e., purification tag). Top left corner states the number of bioreplicates per condition. Data were normalized using in-exchange (0% D) and fully deuterated (100% D) controls; control data points are not shown. Points are means ±SD.

| Precursor MH+ (Da) | Theoretical Mass (Da) | Delta M (ppm) | Peptide 1 | Protein 1 | From | Peptide 2 | Protein 2 | From | Max XlinkX score | Total CSMs | Datasets |
|---|---|---|---|---|---|---|---|---|---|---|---|
| 2466.19 | 2466.19 | -0.03 | ATHSAG[K]R | DCC (cytoplasmic) | 1140 | D[K]GYTTLQDEAIK | M4F | 1560 | 148.89 | 4 | 3 |
| 2037.06 | 2037.06 | -0.93 | ATHSAG[K]R | DCC (cytoplasmic) | 1140 | AYISMIV[K]K | M4F | 2042 | 168.13 | 6 | 4 |
| n/a | n/a | n/a | ATHSAG[K]R | DCC (cytoplasmic) | 1140 | [K]RYSTTR | M4F | 2043 | 101.25 | 1 | 1 |
| n/a | n/a | n/a | ATHSAG[K]R | DCC (cytoplasmic) | 1140 | YALFTYESL[K]K | M4F | 1677 | 130.98 | 1 | 1 |
| 1684.97 | 1684.87 | -0.8 | ATHSAG[K]R | DCC (cytoplasmic) | 1140 | FHL[K]R | M4F | 1654 | 146.61 | 3 | 3 |
| 2775.37 | 2775.37 | 4.29 | ATHSAG[K]R | DCC (cytoplasmic) | 1140 | FE[K]LAATSEVGDLPWK | M4F | 1770 | 94.89 | 1 | 1 |
| 2539.20 | 22539.20 | -0.86 | ATHSAG[K]R | DCC (cytoplasmic) | 1140 | ISQST[K]TFTPCER | M4F | 1869 | 209.7 | 2 | 1 |
| 2851.35 | 2851.35 | -1.43 | ATHSAG[K]R | DCC (cytoplasmic) | 1140 | ENCLNSDVVEQIY[K]R | M4F | 1528 | 145.55 | 1 | 1 |
| 4632.33 | 4632.33 | 0.64 | ATHSAG[K]R | DCC (cytoplasmic) | 1140 | GEGRPLEVFQYEHILSFGAPLANTY[K]IVVDER | M4F | 2011 | 109.38 | 1 | 1 |
| 2685.42 | 2685.42 | 0.67 | [K]GSQK | DCC (cytoplasmic) | 1142 | ITINSHTTAGEVVE[K]LIR | M4F | 1728 | 127.38 | 3 | 3 |
| 2551.29 | 2551.29 | 0.01 | NIE[K]PSGTDPAGR | DCC (cytoplasmic) | 1167 | AYISMIV[K]K | M4F | 2042 | 134.91 | 1 | 1 |
| 3365.58 | 3365.57 | -0.87 | NIE[K]PSGTDPAGR | DCC (cytoplasmic) | 1167 | ENCLNSDVVEQIY[K]R | M4F | 1528 | 193.47 | 1 | 1 |
| 2492.36 | 2492.36 | -0.86 | THV[K]TASLGLAGK | DCC (cytoplasmic) | 1380 | AYISMIV[K]K | M4F | 2042 | 208.17 | 4 | 4 |
| 2564.34 | 2564.34 | -0.11 | TASLGLAG[K]AR | DCC (cytoplasmic) | 1389 | YALFTYESL[K]K | M4F | 1677 | 187.31 | 2 | 2 |
| 2683.36 | 2683.36 | -1.1 | TASLGLAG[K]AR | DCC (cytoplasmic) | 1389 | D[K]GYTTLQDEAIK | M4F | 1560 | 188.87 | 2 | 2 |
| 3182.72 | 3182.72 | 0.96 | TASLGLAG[K]AR | DCC (cytoplasmic) | 1389 | ITINSHTTAGEVVE[K]LIR | M4F | 1728 | 202.79 | 3 | 2 |

**Supplementary Figure 7. Detected cross-links between cytoplasmic DCC and Myo10 M4F.** MS2 details are reported for the peptide spectrum with the highest Max XlinkX score. There are 38 and 15 lysines in M4F and cytoplasmic DCC, respectively. The crosslinked lysine residue numbers correspond to the position in the full-length protein sequences. 5 total datasets analyzed.

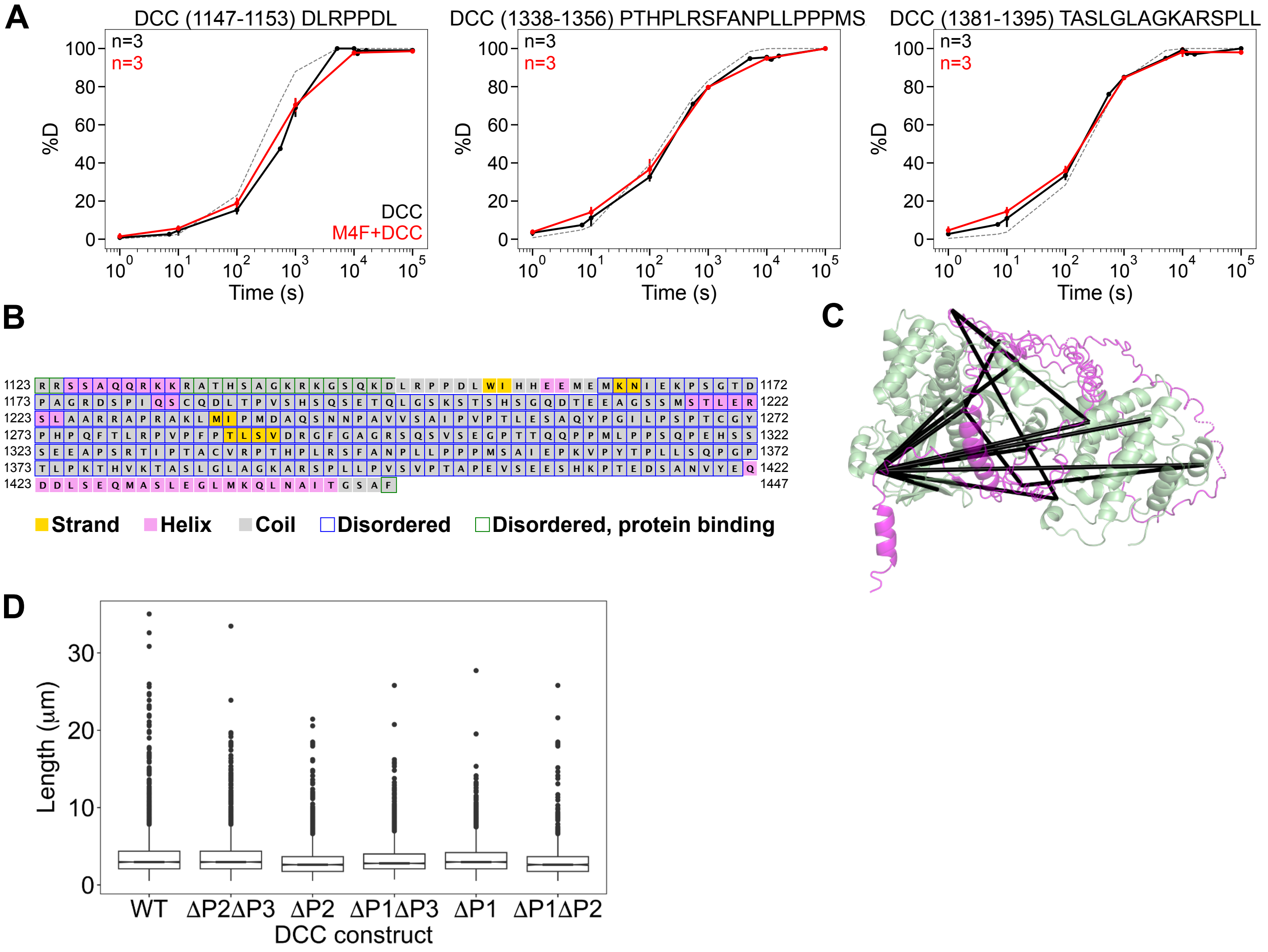


**Supplementary Figure 8. Crosslinking sites in DCC map to true IDRs. DCC truncation mutants do not greatly alter filopodial length in cells.** A) Deuterium uptake plots for peptides covering regions where crosslinks were detected on cytoplasmic DCC. Numbers above each plot correspond to the residue positions on the full-length DCC sequence. Peptide 1147-1153 partially covers the P1, peptide 1338-1356 covers P2, and peptide 1381-1395 is downstream of P2. Unbound DCC = black line, bound to M4F (single-chain construct) = red line, $k_{chem}$ = gray dashed line. Top left corner states the number of bioreplicates per condition. Data were normalized using in-exchange (0% D) and fully deuterated (100% D) controls; control data points are not shown. Points are means ±SD. B) DISOPRED3 annotation of cytoplasmic DCC sequence. C) DSSO crosslinks (black) mapped onto an AlphaFold-Multimer M4F (green) and cytoplasmic DCC (magenta) structure. D) Length of filopodia for the different DCC mutation constructs analyzed in Figure 3D (WT, 2652 filopodia, 90 cells; ΔP2ΔP3, 2966 filopodia, 121 cells; ΔP1ΔP3, 2172 filopodia, 121 cells; ΔP1ΔP2, 1143 filopodia, 93 cells; ΔP1, 2461 filopodia, 98 cells; ΔP2, 1497 filopodia, 91 cells).

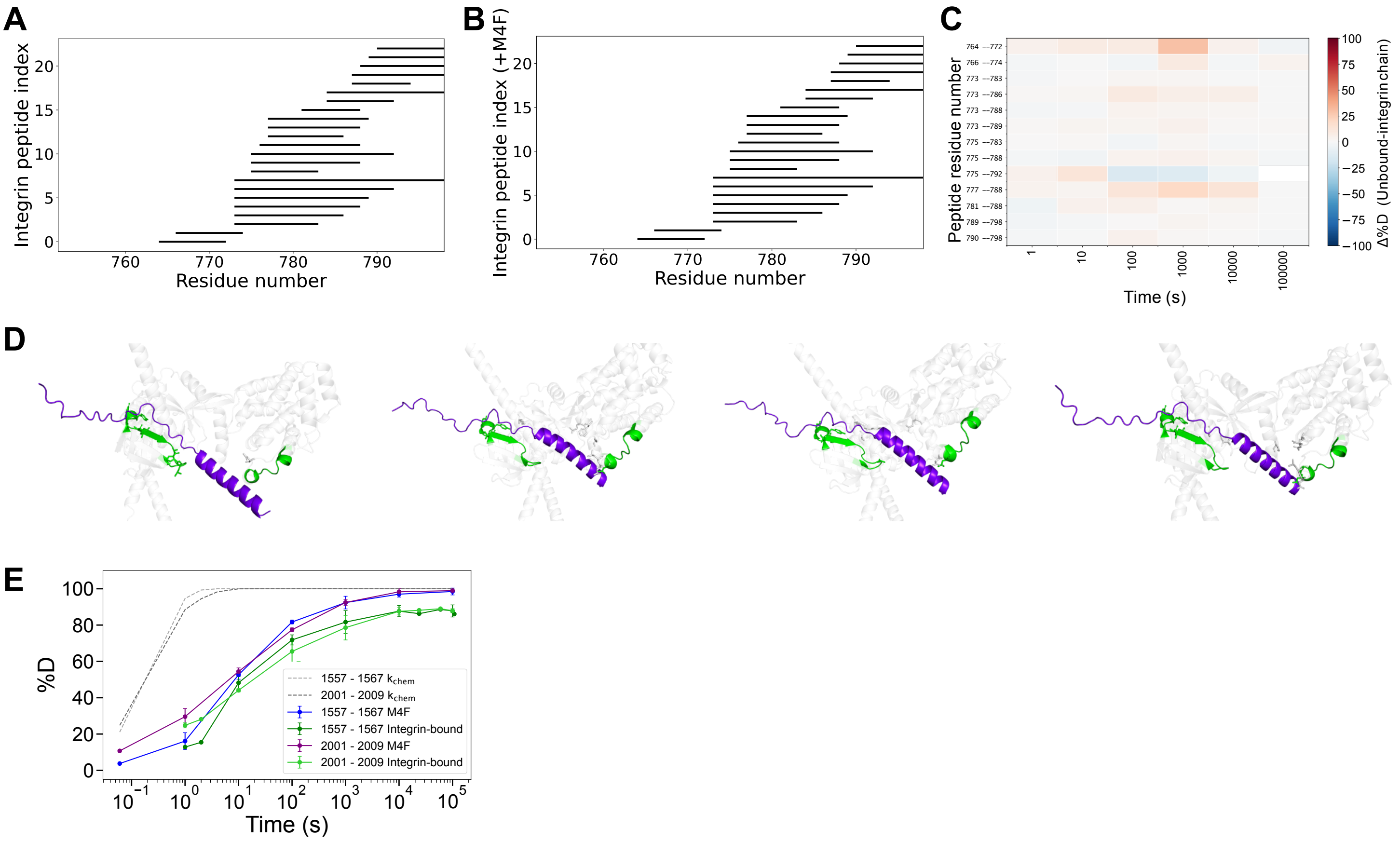


**Supplementary Figure 9. HDX-MS sequence coverage of cytoplasmic β1 integrin and deuteration differences upon Myo10 M4F binding.** Peptide sequence coverage for A) unbound cytoplasmic β1 integrin, B) cytoplasmic β1 integrin bound to M4F (tethered construct). The residue numbers correspond to the position in the full-length protein sequences. C) Heatmap showing deuteration changes for peptides detected in both unbound and M4F-bound cytoplasmic β1 integrin conditions across labeling time points. Red indicates less deuteration upon binding. Peptide residue ranges are displayed on the y-axis. D) AlphaFold3 models of cytoplasmic β1 integrin structure (purple) bound to M4F (light gray). M4F peptides showing HDX-MS stabilization upon binding are in green. M4F residues within 3.2 Angstroms of cytoplasmic β1 integrin are rendered as “sticks.” Proposed models all indicate interactions between cytoplasmic β1 integrin and the M4F peptides that showed protection in the bound complex. E) Overlaid deuterium uptake plots of M4F peptides 1557-1567 and 2001-2009 from Figure 4B.

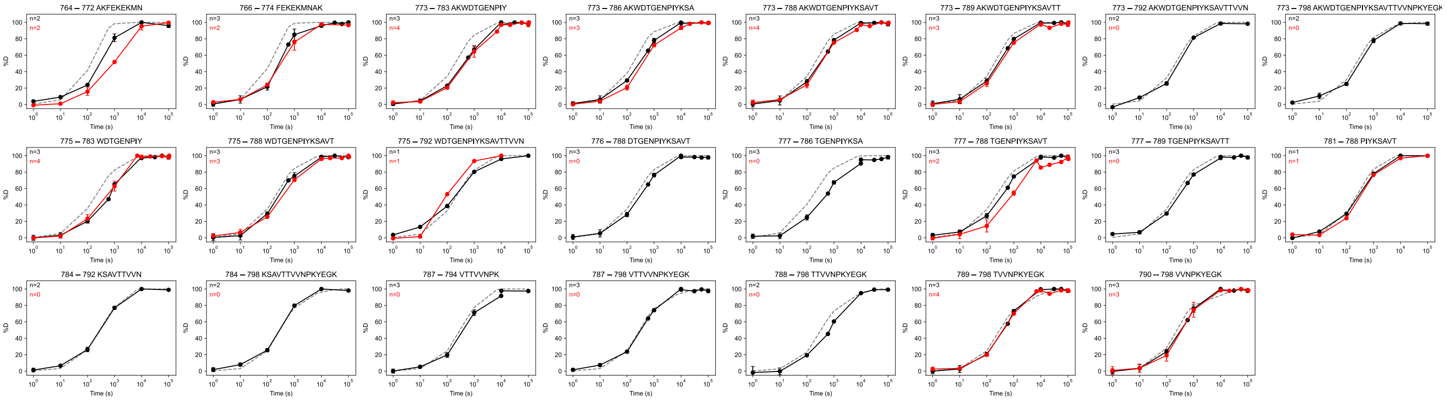


**Supplementary Figure 10. Deuterium uptake plots for all cytoplasmic β1 integrin peptides.** Unbound cytoplasmic β1 integrin = black line, bound to M4F (single-chain construct) = red line, $k_{chem}$ = gray dashed line. Numbers above each plot correspond to the residue positions on the full-length β1 integrin sequence. Top left corner states the number of bioreplicates per condition. Data were normalized using in-exchange (0% D) and fully deuterated (100% D) controls; control data points are not shown. Points are means ±SD.

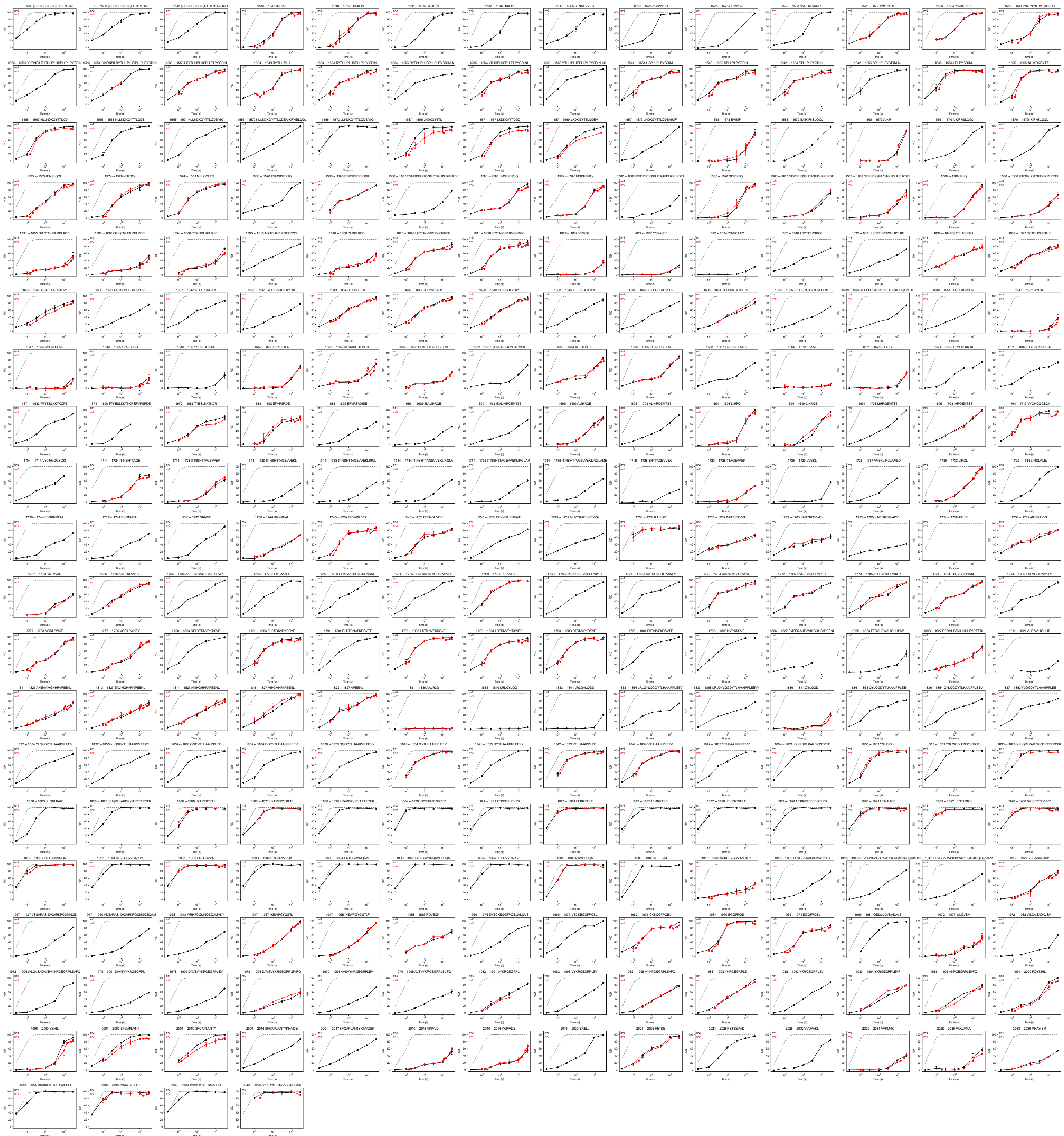

**Supplementary Figure 11. Deuterium uptake plots for all Myo10 M4F peptides.** Unbound M4F = black line, bound to cytoplasmic β1 integrin (single-chain construct) = red line, $k_{chem}$ = gray dashed line. Numbers above each plot correspond to the residue positions on the full-length Myo10 sequence. Grayed numbers do not belong to the protein sequence (i.e., purification tag). Top left corner states the number of bioreplicates per condition. Data were normalized using in-exchange (0% D) and fully deuterated (100% D) controls; control data points are not shown. Points are means ± SD.

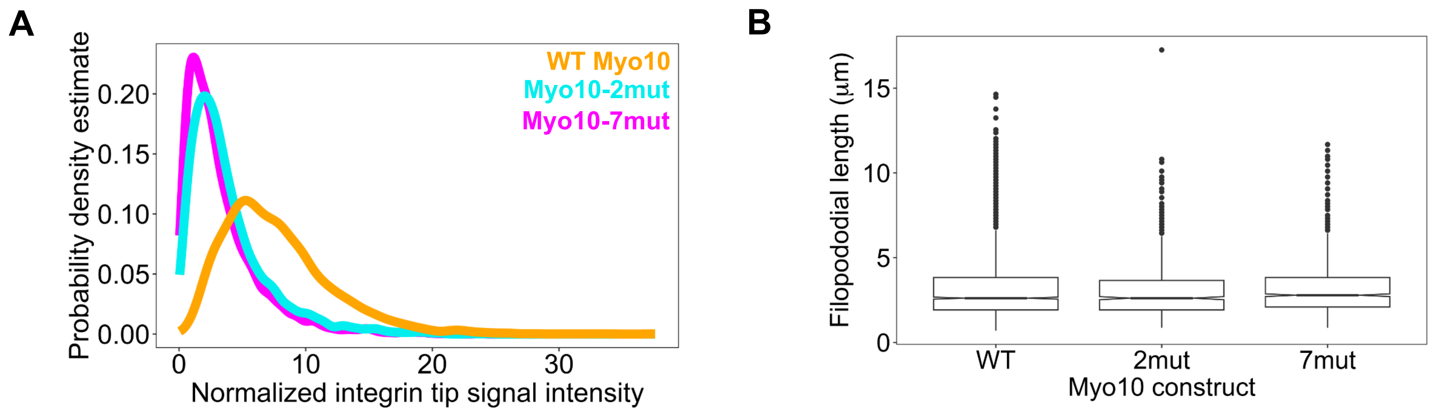


**Supplementary Figure 12. Myo10 mutants that are β1 integrin binding-incompetent are less colocalized with Myo10 at filopodial tips but have similar filopodial lengths in cells.** A) Probability density estimate of the signal intensity of the Myo10 mutation constructs at tips of filopodia with length > 2 μm (WT, 1443 filopodia, 115 cells; 2-mut, 493 filopodia, 42 cells; 7-mut, 687 filopodia, 45 cells). B) Length of filopodia for the different Myo10 mutation constructs analyzed in Figure 4D (WT, 2435 filopodia, 115 cells; 2-mut, 880 filopodia, 42 cells; 7-mut, 1165 filopodia, 45 cells).

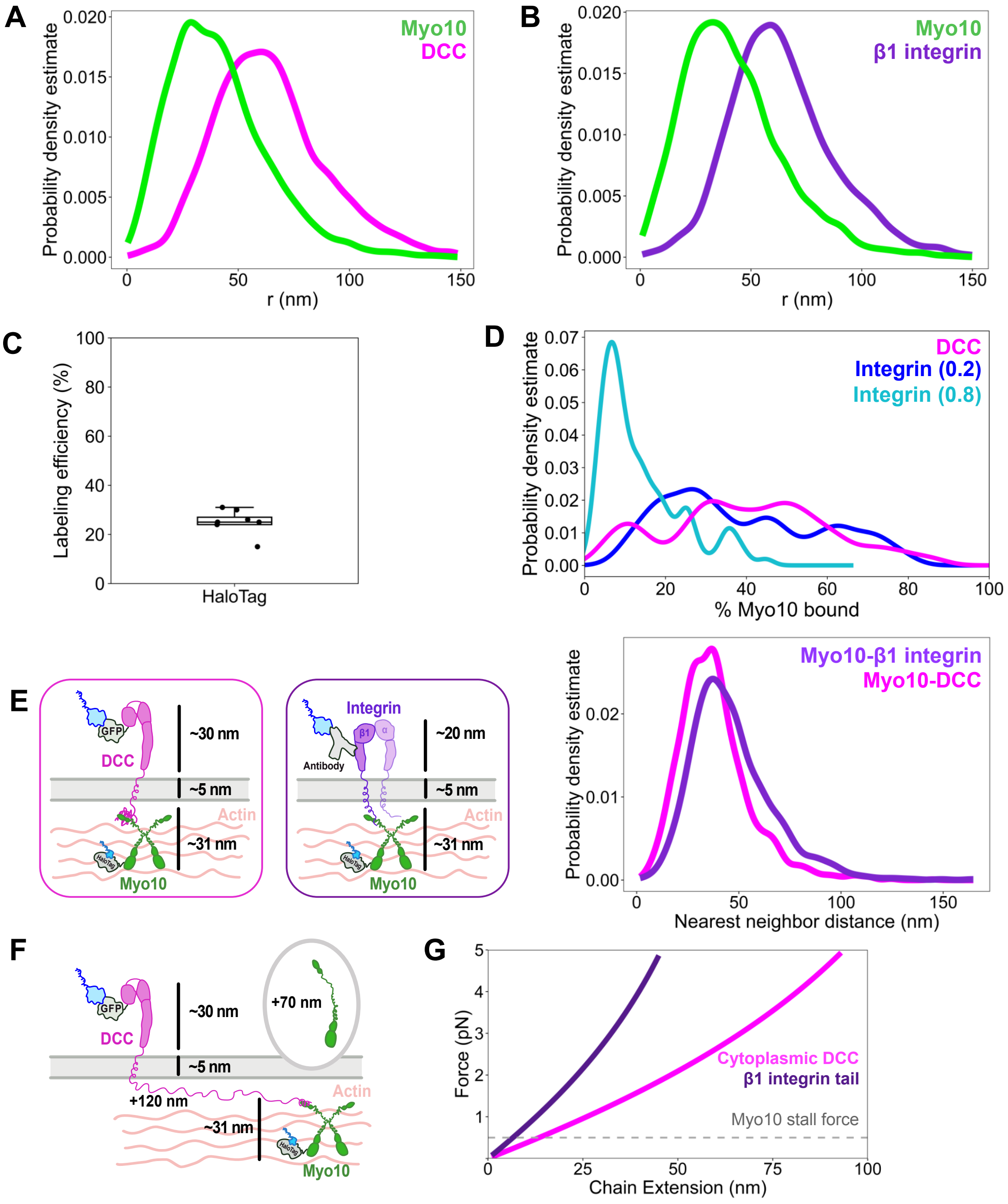


**Supplementary Figure 13. Distances between Myo10 and cargo in filopodia.** A) Polar coordinate mapping of Myo10 and DCC molecule positions in 147 filopodial sections analyzed from 10 cells and B) Myo10 and β1 integrin molecules in 106 filopodial sections analyzed from 7 cells. Molecule positions were projected onto the plane orthogonal to the filopodium's principal axis (PC1) and converted to polar coordinates. Distribution of radius r (in nm) of Myo10 and cargo in filopodia. C) Labeling efficiency of HaloTag probe used for DNA-PAINT. Mean ± SD = 25% ± 4.84% of 8 cells. D) Distribution of % Myo10 bound by cargo, calculated using the quadratic binding equation ($K_D$: DCC = 2 μM; β1 integrin = 25 μM, published). DCC in magenta; integrin in dark blue (20% LE) and cyan (80% LE). E) Left: Models of unloaded Myo10 (not actively exerting force) and cargo (left = DCC, right = β1 integrin) nearest neighbor distances, accounting for the membrane bilayer thickness, protein lengths, and labeling probe docking sites. Right: Distribution of the mutual nearest-neighbor distances between cargo and Myo10 molecule positions; distances < 150 nm are plotted. DCC median = 36.1 nm (n = 1819 distance pairs). β1 integrin median = 43.2 nm (n = 1129 distance pairs). F) Left: Diagram of extended cytoplasmic DCC domain lengths. In a scenario where the cytoplasmic region of DCC is stretched to its chain length, estimating each residue as 0.4 nm, an additional 120 nm of distance would be added to the unloaded Myo10 reference state. If the IDR region of Myo10 (the PEST domain) was extended, an additional 70 nm of distance would be added. G) Force extension curve comparing cargos, where the persistence length of a worm-like chain IDR approximated as 0.8 nm. Cytoplasmic DCC (until the P3 helix, 300 residues), including an extended Myo10 PEST domain (237 residues), is estimated as a chain length of 214.8 nm. Cytoplasmic β1 integrin (estimating 25 residues that are not involved in Myo10 contacts), including an extended Myo10 PEST domain (237 residues), is estimated as a chain length of 104.8 nm. The gray dashed line represents the Myo10 stall force at 0.5 pN. DCC extension at 0.5 pN = 13.48 nm, compliance = 0.038. β1 integrin extension at 0.5 pN = 6.57 nm, compliance = 0. 0.079.
